# Robust and High-Throughput Flickering Spectroscopy for Measurement of Red Blood Cell Membrane Mechanics

**DOI:** 10.64898/2026.09.16.751779

**Authors:** Filip Ayazi, Jurij Kotar, Julian C. Rayner, Pietro Cicuta

**Affiliations:** Cavendish Laboratory, University of Cambridge, Cambridge CB3 0HE, United Kingdom; Cambridge Institute for Medical Research, University of Cambridge, Cambridge CB2 0XY, United Kingdom

## Abstract

The mechanical properties of red blood cell (RBC) membranes are critical to their function in oxygen delivery, and changes to these properties as RBCs age affect their journey around the circulatory system, including clearance by the spleen. Such changes can also have significant health effects, including in cardiovascular disease and on the interactions between RBCs and malaria parasites. The function of RBCs requires them to be very soft, which together with the small size of the cells brings the energy required for measurable deformation of the membrane into the range of typical thermal energies. Red Blood Cells can be observed to flicker under optical microscopy, and these shape fluctuations can be quantified and used to obtain key biophysical parameters such as the tension and bending modulus of the membrane. Typically the shape of the cell’s equator is extracted, and the mean power spectrum is obtained by time-averaging the power present in the normal modes of the thermal fluctuations. Flickering spectroscopy has been used extensively on RBCs, but has so far been very low throughput and with some technical limitations. Here we address issues related to active versus passive fluctuations, focusing and optics, camera exposure and sampling, and fast contour detection. In combination with an automated imaging system it is possible to measure thousands of cells in one day, with no user input. We validate this new pipeline by chemical modification of RBC mechanics, and by comparison with simulated fluctuations including each confounding effect. The methods and codes for this robust flickering analysis will allow consistent measurements across labs.

**SIGNIFICANCE:** Mechanical properties of red blood cell (RBC) membranes are critical to their function and are altered in a range of conditions, yet measuring membrane mechanics cell by cell remains slow and sensitive to analysis choices. Flickering spectroscopy measures membrane mechanical properties from thermal fluctuations of the membrane, but suffers from low-throughput, large labour requirements and technical limitations due to non-thermal effects and instrumental artifacts. We present an automated imaging and analysis pipeline that allows the measurement of thousands of cells per day without user input and corrects for the artifacts. Validated on simulated fluctuations and chemically stiffened cells, this new pipeline gives robust measurements with the throughput and automation needed to run large-scale studies and compound screenings.

## 1 INTRODUCTION

The mechanical properties of red blood cells (RBC) are critical to their function in oxygen delivery, as the cells repeatedly deform in circulation passing through narrow capillaries. Changes to these mechanical properties can have significant health effects and they serve as a mechanism for RBC ‘quality control’: as cells age, their mechanical properties change (1, 2), and this loss of deformability leads directly to their clearance by the spleen. These mechanical properties are altered in conditions affecting cell shape (3, 4), but also in many other conditions (5–12). Given that the interior of the RBC is normally a fluid, the mechanics of the cell are dominated by the membrane. The membrane, as a thin sheet, is often considered to have a resistance to stretch (tension), to bending (bending modulus), and to shear (shear modulus). Even small changes in cell tension can have significant effects, for example we previously identified a protective role of high tension against malaria parasite invasion, in a rare human genetic trait (13). The importance of these parameters and the range of diseases involved highlights the need for an effective and scalable method of their measurement.

Because the function of RBCs requires them to be very soft, and given their small size, the energy required for measurable deformation of the membrane is on the order of typical thermal energies. As a result, the cells are seen to continuously fluctuate shape (or “flicker”) under optical microscopy. This effect has been known for well over a century (14). With advances in theoretical models of the membrane (15, 16), and in imaging and digital processing technologies, more quantitative studies became possible over time. The experimental technique shifted from recording the temporal spectrum of the intensity fluctuations at a point in the cell, to measuring the fluctuations of the shape of the cell, mainly focusing on the shape of the ‘equator’ (15, 17–19), see Figure 1. This technique of studying RBC has benefited from the parallel developments of flickering spectroscopy as a method of studying fluctuating vesicles. Lipid vesicles are typically spherical, and model systems can be made with well controlled properties, leading to technical advances in this field, notably models by (20, 21). In particular two papers are foundational: (1) Pécréaux et al. (20) developed a technique for measuring the tension and bending modulus on giant unilamellar vesicles based on flickering, under the assumption of zero depth of field focusing on the observed equator, and have shown that this simplified 2D model is equivalent to a spherical model at high enough spatial modes. That work also properly considered the effect of camera exposure times. These methods, developed for spherical vesicles, have been successfully used on RBCs (which are not spherical) by us and others (13, 22–24) because the effect of the different radius of curvature is only present at long wavelengths which are not used in analysis. Using this technique, and assuming a thermal nature of the fluctuations, it is possible to extract membrane tension *σ*, bending modulus *κ*, and by using the temporal relaxation dynamics of excitations it is also possible to extract the effective viscosity *η*, which is typically assumed to be dominated by the internal viscosity of the cell (22). The second foundational paper is (2) Rautu et al. (21), in which the treatment of (20) was extended to account for the finite depth of field of actual microscope setups, testing on spherical vesicles.

**Figure 1:**
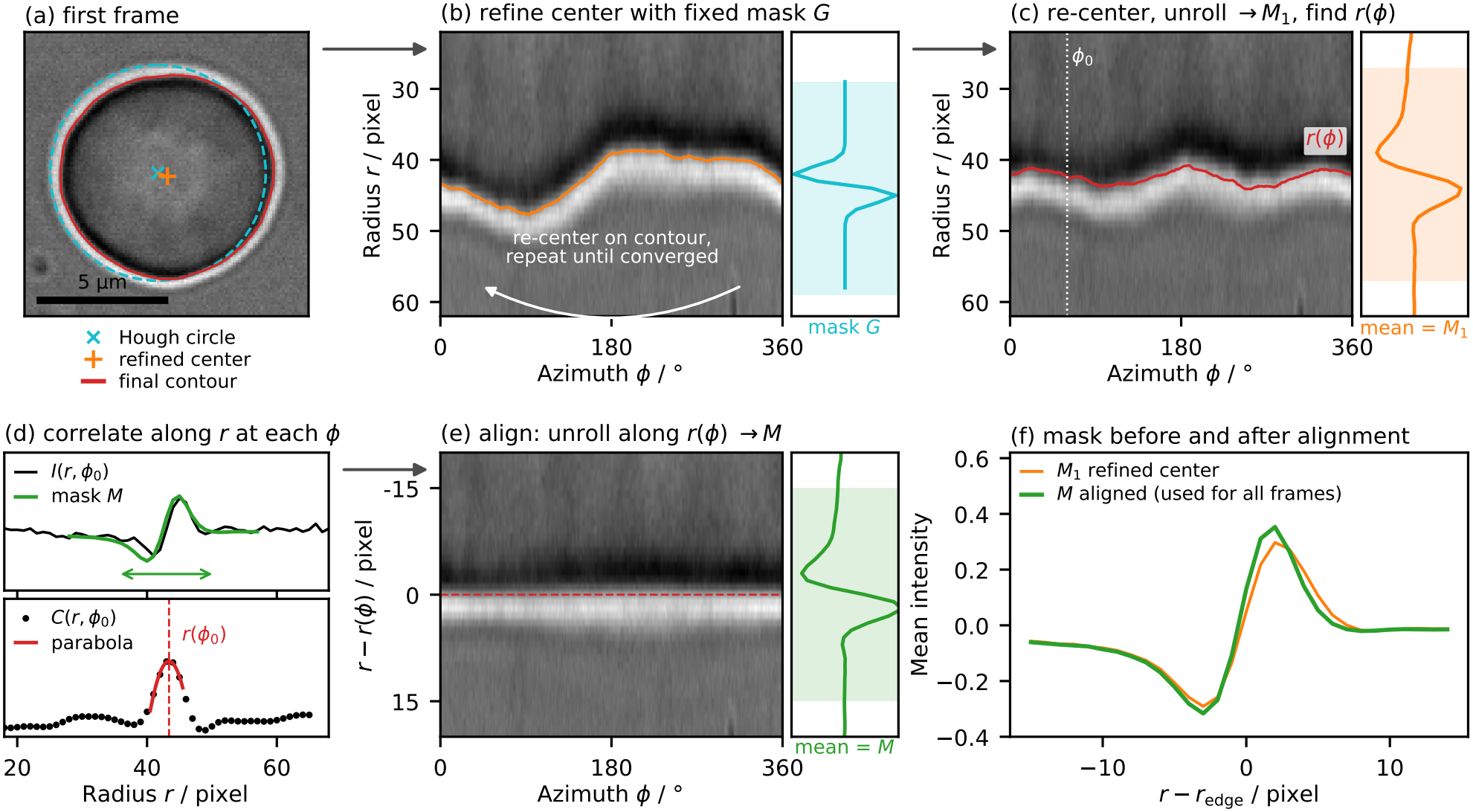
Workflow for correlation-based contour detection. (a) First frame of a video, with the initial estimate of the cell center and membrane position from a Hough transform (cyan), the refined center (orange) and the final detected contour (red). (b) The image is unrolled around the estimated center, i.e. resampled along radial lines to give the intensity *I* (*r, ϕ*) as a function of radius *r* and azimuth *ϕ*, and each radial line is correlated with a predefined mask *G* based on the gradient of a Gaussian. The position of maximum correlation, located to sub-pixel precision by fitting a parabola around the peak (d), gives an estimate of the contour *r* (*ϕ*). The center of mass of this contour is used as the new center estimate, and the procedure is repeated until the center converges. (c) The image is unrolled around this refined center and averaged across azimuths to give an image-derived mask *M*_1_, which is correlated with the unrolled image as before to obtain a new estimate of the contour. For non-circular contours the membrane lies at different radii at different azimuths, so *M*_1_ is blurred. (e) The image is therefore unrolled around the contour, shifting each azimuth so that the contour becomes a straight line, and averaging across azimuths gives the mask *M*, which best represents the radial intensity profile of the membrane; (f) compares *M*_1_ and *M*. The mask *M* is then fixed and used for correlation-based detection on all frames of the video, including the center-finding iteration, with each contour refined by a second correlation pass on the image unrolled around it. Using an image-derived mask makes the method more robust to slight defocus than a predefined mask. On an Intel Core i7-14700K CPU for typical videos the first-frame processing and mask extraction take approximately 44 ms. Once the mask is extracted and initial center estimated, subsequent frames are processed in parallel, taking approximately 5.1 ms per frame on a single core and running on all cores the system processes on average 914 frames per second.

Here we combine these various technical aspects to the study of red blood cells, specifically improving the whole pipeline of flickering experiments and analysis to enable high-throughput, automated flickering spectroscopy of RBCs for the first time. We present a fully automated optical microscopy system capable of imaging thousands of RBCs per day, significantly reducing manual labor and input, and increasing robustness and statistical power. We also develop an analysis pipeline using frequency filtering to separate thermal fluctuations from non-thermal effects, and we improve mode dynamics analysis and exposure time correction by making decay time calculation more resistant to the non-thermal contributions. We validate our approach with simulations and by treating the cells with glutaraldehyde which is known to affect membrane tension.

## 2 BACKGROUND ON TECHNICAL ASPECTS OF FLICKERING

### 2.1 Spherical and Equatorial modes

The modern approach to flickering spectroscopy works by extracting the fluctuating shape of the cell’s equator observed and segmented over time from optical microscopy images, and then obtaining the mean power spectrum of the normal modes. From this spectrum one can determine the tension (*σ*) and bending modulus (*κ*) of the membrane (15).

Most treatments are primarily based on a flat membrane model, which has been shown to be equivalent to a spherical model at higher modes. In this model, the mean power in modes with wavenumber *q* on the equator of a cell is given by (20)

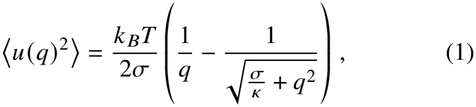

where *u* (*q*) is the complex amplitude of the mode with wavenumber *q*. A number of technical/experimental considerations are essential for performing this analysis and are described below.

### 2.2 Accounting for exposure time

In practice, the finite exposure time of the camera (*T*_exp_) leads to an observed attenuation of the fluctuation power. For a thermal mode with a decay time *τ*, the observed power spectrum is given by (20):

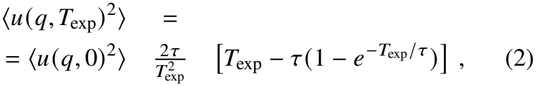

where *⟨u* (*q*, 0)^2^⟩ is the true fluctuation power. This takes account of the averaging out of fast modes.

### 2.3 Accounting for optical focus depth

The technique aims to record the one-dimensional position of the cell’s equator, but experimentally this is not possible: The image is formed averaging over a (thin but not negligible) thickness. This means that the simple assumption leading to (20) needs to be challenged, because the recorded equator contains signal from perpendicular fluctuation modes. This was done by Rautu et al. (21), accounting for the depth of field of a microscope when imaging spherical vesicles. While the surface of a red blood cell is not spherical, the observed part of the cell’s equator is close to the edge, and a spherical approximation is reasonable for the wavelengths of interest.

In this spherical approach the fluctuations are expressed using spherical harmonics 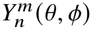. The mean-squared amplitudes of these true 3D shape fluctuations, 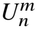, are described by the Helfrich Hamiltonian (25):

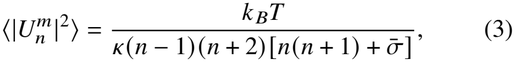

where *k* _*B*_*T* is the thermal energy, *κ* is the membrane bending modulus, *n* ≥ 2 and |*m*| ≤ *n* are the integer degree and order of the spherical harmonic, and 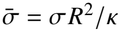 is the dimensionless reduced surface tension.

When analyzing the RBC contour via optical microscopy, the finite focal depth of the objective causes out-of-focus membrane fluctuations near the cell’s equator to be projected onto the 2D imaging plane. To correct for this spatial averaging, the acquired optical signal is modeled as a convolution of the membrane shape with a Gaussian kernel whose width depends on the focal depth.

The procedure above is rigorous in fluorescence microscopy. By defining the dimensionless focal depth parameter Δ as the ratio of the microscope’s focal depth to the mean equivalent sphere radius *R*, the experimentally observable 1D Fourier modes along the RBC equator, *⟨*|*µ*_*q*_ | ^2^ ⟩, can be expressed as a weighted sum of the 3D spherical modes:

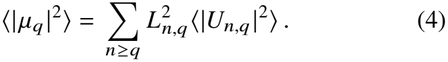

In practice what is measured from videos of red blood cells are the radial positions of the membrane *r* (*ϕ* _*j*_, *t*) at *N* equally spaced azimuths *ϕ* _*j*_ = 2*π j/N*. To relate this to the quantities derived here we convert these into the Fourier modes 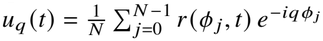 (with the inverse transform 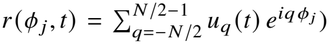, whose *q* = 0 mode gives the mean radius *R*. These are related to the dimensionless modes above by *u*_*q*_ = *Rµ*_*q*_, so the measured spectrum is fitted with *⟨*|*u*_*q*_| ^2^⟩ = *R*^2^ *⟨*|*µ*_*q*_ |^2^⟩ using Eq. 4.

The optical projection coefficients, *L*_*n,q*_, correct for the spatial averaging of the out-of-focus RBC membrane. These coefficients are analytically determined as:(21)

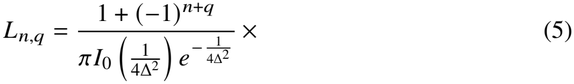

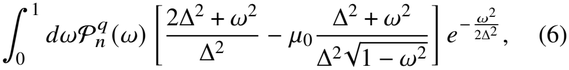

where 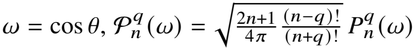 are the normalised associated Legendre polynomials, with 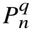 the associated Legendre polynomials (so that 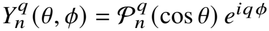 and *I*_0_ is the modified Bessel function of the first kind of order zero. The term 1 + (−1)^*n*+*q*^ represents the factor of 2 coming from the restricted limit of the integration over the even function.

The factor *µ*_0_ accounts for the zeroth-order term in the expansion of the projected radius and is evaluated as:(21)

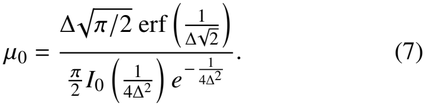

In the ideal limit of an infinitely thin focal depth (Δ →0), *µ*_0_ →1 and the projection coefficients reduce to the standard analytical values for an exact equatorial slice, eliminating the out-of-focus contributions (21). We show later in Section 4.2 how this is applied to RBCs in brightfield imaging.

### 2.4 Thermal and non-thermal fluctuations

The models discussed above describe fluctuations in thermal equilibrium, where the amplitude of each mode is set by *k* _*B*_*T* and the membrane properties. Observed fluctuations are often also affected by other processes such as cell rotation or flows in the chamber, which are generally slower than the fast timescale of purely thermal fluctuations. Apart from these experimental effects, recent discussion has focused on whether the observed RBC shape fluctuations are entirely thermal, or if there is an active (non-equilibrium) contribution arising from biochemical processes. While some authors have reported no ATP dependence, indicating a purely thermal nature of the fluctuations (24, 26–28), others have observed ATP dependence (29–34), and data inconsistent with thermal fluctuations only (35). The observations of active fluctuations generally consistently show the active component to be significantly slower than the thermal contributions - with thermal behaviour dominating at timescales below 100 ms (29, 34–36). These additional effects are not modelled by the equilibrium models and fitting with these in the spectra would bias the parameters and worsen the fit (as seen later in Section 2.6).

We show later in Section 4.3 how this separation of timescales allows us to separate the two types of shape deformations. If active processes were to change the mechanical state of the membrane and cytoskeleton, rather than just adding power to some fluctuations directly, these would be retained through their effect on the measured membrane properties, so the parameters we report describe the fast fluctuations of the cell in its current state. Connecting the mechanical effect of tension obtained from shape fluctuations to particular mechanical deformations is not straightforward: the tension governing slower deformations would be particularly affected by active processes, which can take place on those timescales. Active forces fast and frequent enough would also couple into the thermal modes, adding to their driving forces, and acting as an increased effective temperature in what would appear like a Helfrich fluctuation spectrum. Such effective temperature would rescale both tension and bending modulus (30), even though the direct measurements find no such excess at timescales below 100 ms (35). So the interpretation of absolute tension values remains complex, but identifying and removing “non-Helfrich” contributions by filtering allows a standardisation of the analysis, and opens the way to reliably detect changes in membrane tension. In conclusion here, we anticipate that in our experiments testing the membrane fluctuations for time asymmetry (37) and other metrics of non-equilibrium (34) has not shown evidence of non-equilibrium effects, so in these RBC in our conditions we can say that once non-thermal slow deformations are filtered, the remaining fluctuations are Helfrich-like (by construction of the filter) and also thermal.

### 2.5 Previous pipelines have been low throughput

Despite the considerable power and flexibility of flickering spectroscopy, in practice its use is limited by several considerations. There is a large natural (physiological) variance of membrane properties within a population of circulating RBCs even from an individual donor. For example, cells become stiffer over their c. 120 day lifetime. Hence a large number of cells must be analyzed for each sample to achieve sufficient precision and detect small effects from drugs or across donors or other conditions. Traditionally, experiments were performed manually: an experimenter would move the microscope stage to find a cell, adjust focus and camera recording area, record a video, and repeat with a new cell. This makes it hard to run the experiments with large number of samples or over long time periods, limiting the amount of data available. The manual nature of this process also makes it nearly impossible to track individual cells over long periods of time, and can potentially introduce selection bias (towards what the experimenter deems ‘good’ cells, on the spot) in a way that is hard to quantify.

### 2.6 Choice of fit range significantly affects obtained parameters

The radial equatorial displacements, at equally spaced angles, are Fourier transformed, giving a series of mode amplitudes. When processing the obtained mean power spectra, the choice of which modes should be used for the fit has never been clearly answered.

At low modes (long wavelengths) the planar approximation, and for non-spherocyte RBCs also the spherical approximation, break down. However, at high modes the behaviour is dominated by bending modulus, so if one wants to recover a value of tension, a choice needs to be made to include modes which are low enough to see the effect of tension, while still high enough to avoid the geometric issues. This is not trivial. The high end cutoff of modes also needs consideration.

The exposure time, resolution of the camera and noise from contour tracking and camera pixel fluctuations all contribute to limiting the maximum mode that should be included.

It is not trivial to choose the optimal range of modes, to best use the available data while avoiding bias from these factors. This results in different authors making different choices, which does have a significant effect on the obtained parameters. Typically it is assumed that by using the same parameters and only comparing the results within the same analysis pipeline, data can be compared internally (13, 38). However, these arbitrary choices make it hard to compare results between groups and introduce a level of uncertainty about the method. If the model correctly explains the behavior, the change of fit range should ideally affect the uncertainty but not the value of the parameters.

## 3 MATERIALS AND METHODS

### 3.1 Automated cell imaging

To address the labor requirements of flickering experiments and increase the amount of available data, we developed a fully automated imaging system based on a Nikon Eclipse Ti-E inverted microscope with a Nikon Plan Apo 60×, 1.4 NA oil immersion objective, a FLIR GS3-U3-23S6M camera, and a custom 655 nm LED light source synchronized to the camera. This wavelength was chosen to minimize phototoxicity. During imaging the samples and objective were heated to 37 °C using custom heaters.

We control the microscope using an in-house “*Temika*” control software which receives commands over a TCP/IP connection from our Python-based software. This control system includes retrieving and transmitting images from the camera, allowing fully automated control with feedback from the camera. This allows us to find suitable cells, run automatic focusing and record cells with no input from the user. This imaging pipeline is summarized in Figure 2.

**Figure 2:**
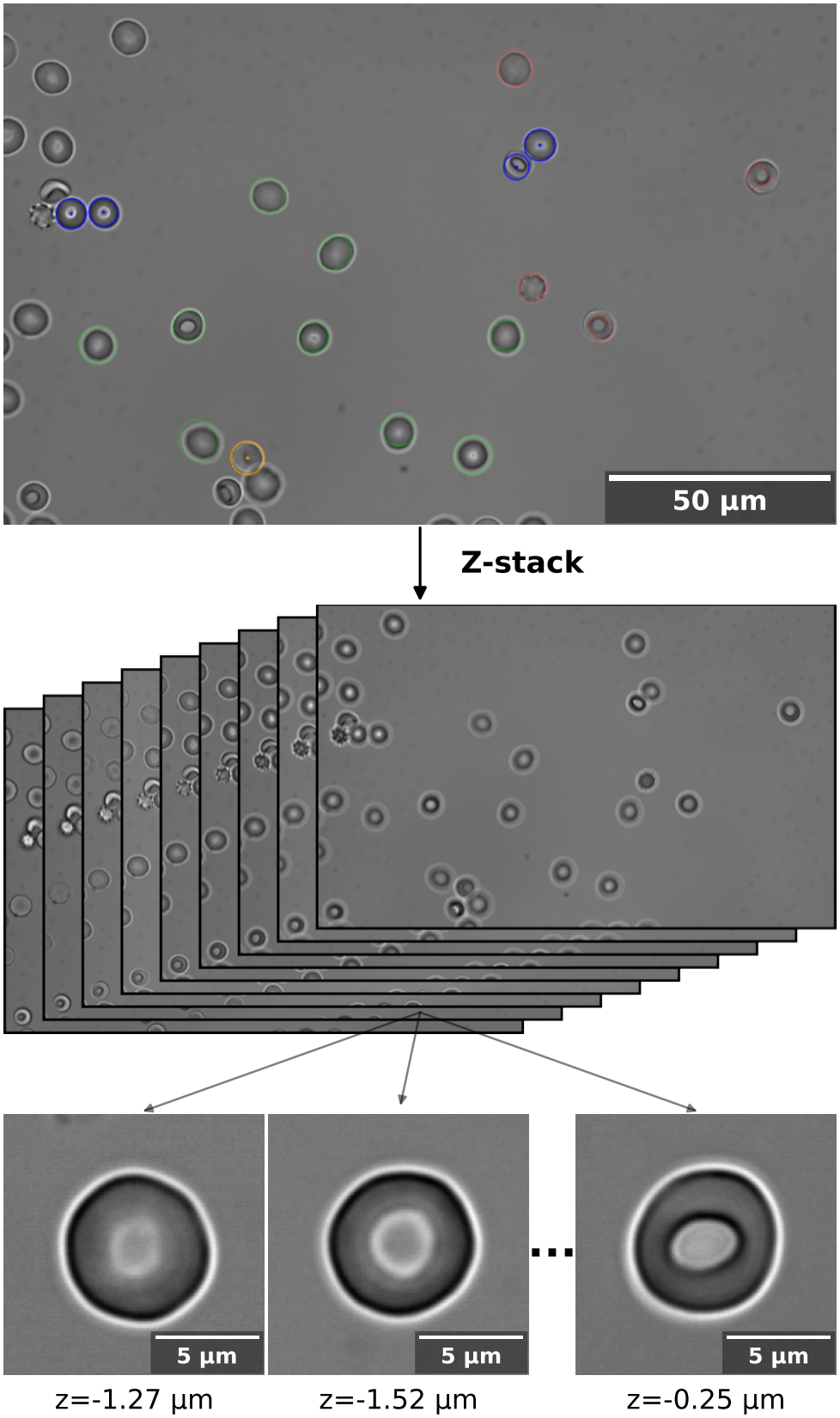
Automatic image acquisition pipeline. The initial evaluation of the full field of view is shown on the top. Cells highlighted in orange were labelled as too deformed or did not have a clear area around them; cells in blue are considered too close to other cells; red outline shows cell contours considered invalid (jagged shape, not a closed contour, high eccentricity); cells in green progress to further processing. A z-stack is obtained to find best focal plane for each cell. For each cell the focus is adjusted to the determined best position, and camera region of interest is then restricted to image each cell at high frame rate.

With this automated system we can image approximately 3500 cells per day with 20 s recordings and approximately 6000 cells per day with 10 s recordings.

#### 3.1.1 Establishing focus requirement

To investigate the focus precision requirements for an automated focus system we used the automated pipeline to record 189 cells at 15 z-positions centered on a plane where the equator appears sharpest (defined as when the mean of maxima of radial gradients in the intensities is maximized). The z-step between recordings was approximately 240 nm. The results of these measurements are presented in Figure 3 and discussed in Section 4.1.

**Figure 3:**
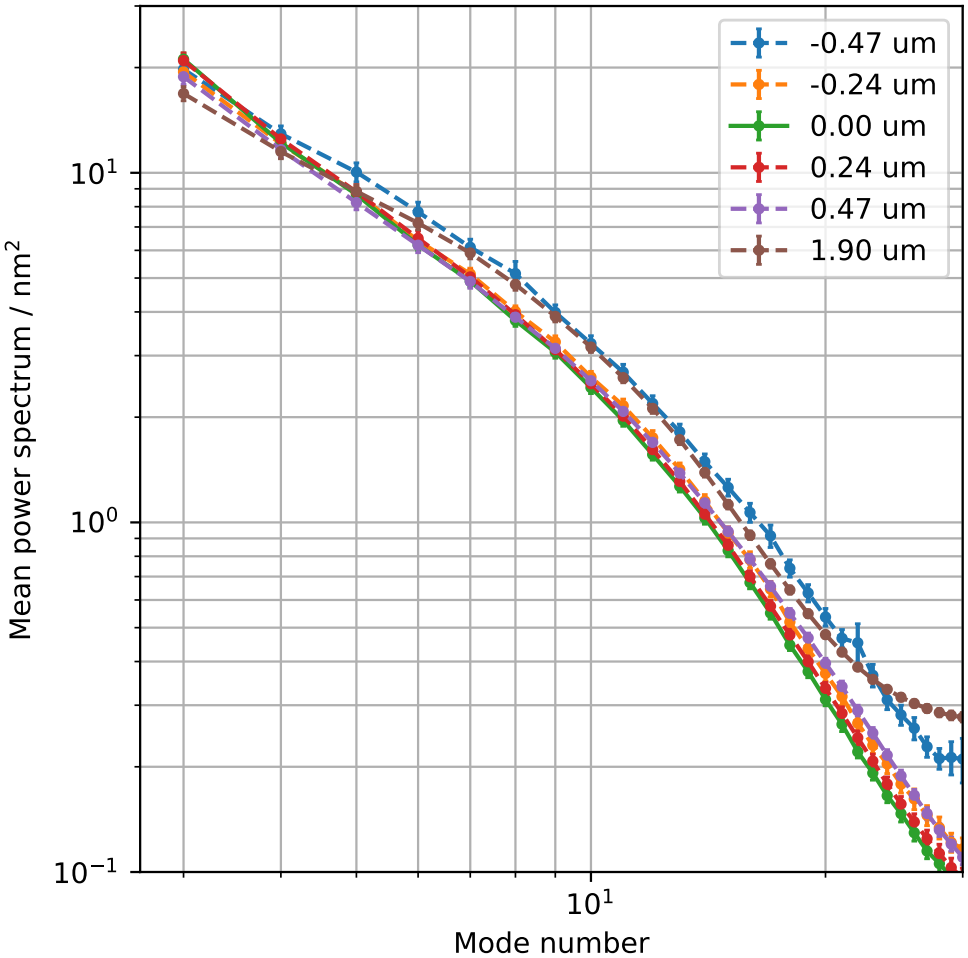
Average power spectrum of thermal fluctuations across a population of red blood cells (*N* = 189) imaged at different focal planes. As the plane goes further from the best-focus plane, the amount of noise increases. Defocusing to a focal plane below the equator appears to result in faster deterioration than planes above. These results are discussed in detail in Section 4.1. The fitted tension and bending modulus as a function of the focal plane position are shown in Figure S5.

#### 3.1.2 Different exposure time imaging

To obtain data on the effect of camera exposure time and study the measurement of fluctuation decay times, we imaged 863 cells at 12 different exposure times between 0.9 ms and 9.9 ms. The shortest exposure time recordings were taken at 660 fps while the rest were obtained at 100 fps. These exposure-time measurements are shown in Figures 11 and 12, and discussed in detail in Section 4.4.

### 3.2 Correlation-based contour detection

The shape of the cell equator needs to be extracted with sub-pixel precision on the radial coordinate. We use a correlation based approach similar to (22), as shown in Figure 1.

To optimize the selected correlation mask, and the width of a parabolic fit within the correlations, interpolated masks were artificially shifted and the accuracy of the shift recovery was tested, giving mask width of 30 pixels and fit width as 5 pixels as the best option. To determine the precision of this method, static images with added noise were tracked, with 10 frames from 16 cells used each used to generate 1000 images by adding normally distributed random noise in different amounts. The results showed recovery of contour position and variance in the static contour to be in single digit nm range for typical noise levels in our videos, and that the expected errors do increase significantly with noise, highlighting the need for low noise videos.

### 3.3 Simulations of equatorial fluctuations

A custom quantitative simulation tool was built to simulate excitations on a line corresponding to the cell equator, simulating overdamped fluctuations on a 1D string representing the cell equator.

At each internal simulation time step of *δt* = 20 µs, new random thermal excitations are added for each mode *n* up to a maximum mode *M* = 50. The phase is randomly and uniformly sampled from [0, *π*]. The initial amplitude of the excitation at its generation time *t*_0_ is drawn from a normal distribution with zero mean and standard deviation:

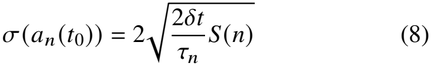

where *S* (*n*) is the expected equilibrium power spectrum (per Eq. 1, or set to a constant) and *τ*_*n*_ is the characteristic decay time of the mode. The amplitude of each excitation relaxes (decays) exponentially:

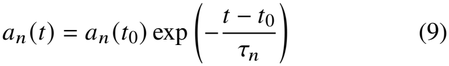

At a set framerate, the total equatorial displacement *u* (*x, t*) is calculated as the sum of the displacements generated by all excitations present at that time. If an excitation amplitude falls below a threshold (set to 5 × 10^−13^ m), it is removed from the simulation.

The decay times used in the simulations (1 ms to 100 ms) were chosen to cover the range expected for RBCs with a large margin on the slow decay side. The relaxation time of a mode depends not only on the viscosity but also on the membrane mechanics, for a typical cell (*R* = 4 µm, *σ* = 15 × 10^−7^ N/m, *κ* = 250 × 10^−21^ J, *η*_in_ = 20 mPa s) using Eq. 13 yields a range of decay times for modes 20-6 *τ* ≈ 1 ms to 13 ms, consistent with our measurements of real cells (Figure 12). The simulated decay times thus cover this range and go up to near 10x the slowest expected thermal mode.

To allow simulations of non-thermal effects, this simulator supports adding additional active excitations which remain constant for a set lifetime before decaying as a thermal mode. In experiments where this was enabled, this lifetime was set to 0.1 s to match reported minimum timescales of active effects (30, 35).

### 3.4 Standard RBC sample preparation and imaging chambers

Red blood cells were purchased from NHS Blood and Transfusion, and ethical approval for their use in this work provided by NHS REC (20/EE/0100). Cells were treated via standard protocols: They were first washed twice with PBS supplemented with 1 mg/ml BSA and 5 mmol/l glucose (wash medium), centrifuging at 3000 RPM between washes. The sample was then diluted to a 0.013 % hematocrit and placed in a hybridization chamber (GBL621501, Grace-Biolabs, US) for imaging.

### 3.5 Treatment of RBCs with glutaraldehyde for validation

Red blood cells were washed twice with PBS supplemented with 1 mg/ml BSA and 20 mmol/l glucose (wash medium), centrifuging at 14 000 RPM between washes. 1 ml of different concentrations of glutaraldehyde in wash medium were prepared and 25 µl of washed RBCs were added to each concentration in rapid succession. The solution was incubated for 30 min at room temperature with moderate shaking. After incubation, all samples were centrifuged at 14 000 RPM for 3 min, supernatant was removed and replaced with 1 ml of wash medium. This washing was repeated 2 more times. After the last washing, supernatant was removed and RBCs were diluted to a final concentration of 1:6700 in wash medium and placed in an acrylic chamber plate.

### 3.6 Custom imaging chambers

Our aim is to improve the flickering pipeline not just in the overall throughput of cell number analysed (which is key to measure distributions of properties, and detect shifts of those distributions with high sensitivity) but also to be able to screen a large number of samples or sample conditions.

When imaging in commercial multiwell plates, we observed significant evaporation, even in a humidified chamber. Furthermore, due to the high walls of each well, the numerical aperture of illumination was limited, reducing both the available light and the resolution in the resulting image. To address both of these challenges we developed a custom imaging plate.

This plate was formed by a stack of a cover glass bottom (GPD-1577, UQG Optics, UK), adhesive, laser cut 1.5 mm thick acrylic spacer with 28 12 mm cutouts forming the chambers, adhesive, and an acrylic lid with 2 1.5 mm fill holes in the corners of each chamber. Adhesive used was 3M 468MP (113317, Self Adhesive Supplies, UK) with cast acrylic (082-4480, RS Components, UK). The chambers were sealed using adhesive dots formed using the same adhesive on a Mylar sheet (7850782, RS Components, UK). The acrylic spacer and lid are cut to the shape of a standard well plate. A diagram representation of these plates is shown in Figure 4. The concept of a laser cut acrylic sandwich chamber was inspired by Kals et al. (39).

**Figure 4:**
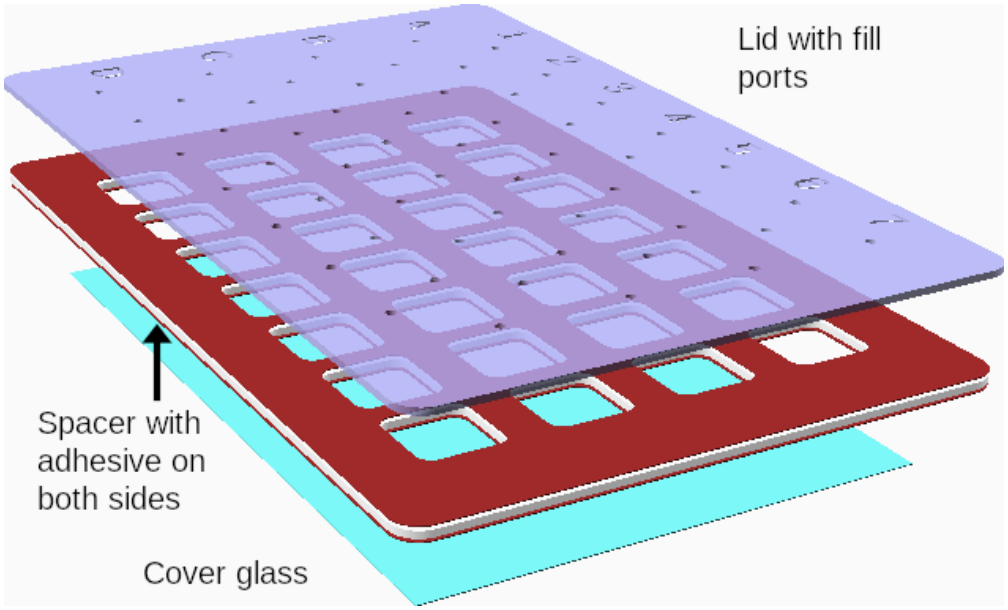
Custom sample chambers facilitate high throughput imaging. Schematic of laser-cut acrylic chambers. The acrylic lid and spacer are laser cut and glued together with a cover slip on the bottom to form chambers, which are sealed with adhesive dots after filling. The plate has the overall dimensions of a standard well plate and is approximately 3.5 mm tall.

In experiments with only a single sample, a commercial hybridization chamber (GBL621501, Grace-Biolabs, US) was used.

The custom design, like the commercial chamber, allows the samples to be sealed and provides a flat top surface which does not interfere with illumination for high resolution imaging. As a significant improvement to the hybridization chamber, this allows us to image a large number of samples in the same experimental run.

### 3.7 Testing and developing methods

We have explained so far the key technical aspects of flickering - these will be tested and validated progressively, to finally create a comprehensive pipeline which is presented in Section 4.7.

## 4 RESULTS

### 4.1 Sensitivity to focal plane position

While the microscope is able to maintain position relative to the cover slip using the Nikon “Perfect Focus System”, as the red blood cell sizes vary so does the height of their equator from the cover slip. This requires adjusting the objective z-position for each cell. Averaging power spectra across the z-stacks of 189 cells, as shown in Figure 3, shows that small changes in focal plane can significantly affect the obtained spectrum. The focus position needs to be within approximately 300 nm of the equator position. In the automated system this is achieved by obtaining an image every 300 nm and fitting a parabola on the focus metric value to obtain the best focus position between the steps, then positioning the focus at this height. The effect of focal plane position on the resulting fit parameters is shown in Figure S5.

### 4.2 Sensitivity to the focal depth correction

In the spectrum analysis, setting the value of Δ, introduced in Section 2.3, has a large effect on the tension and bending modulus obtained from the fits. This is shown in Figure 5.

**Figure 5:**
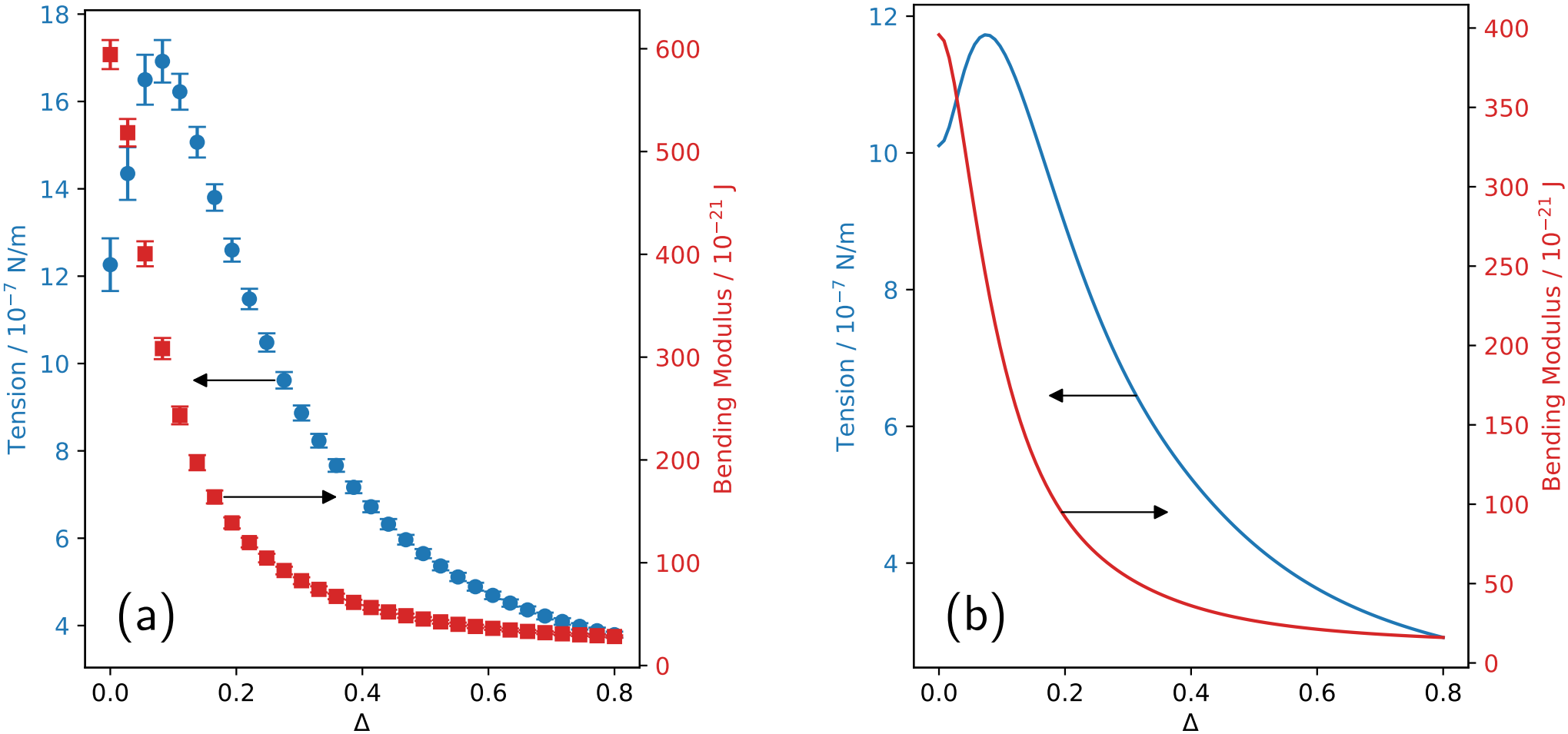
Simulation of fitting with different depth of field parameters Δ on a theoretical spectrum (b) reproduces observed effect on real cells (a). Effect of the depth of field parameter Δ on the obtained mean tension and mean bending modulus calculated across 80 cells, with the error bars showing standard error (a) and the same effect visible in (b) when fitting a theoretical, noise free spectrum generated with Δ = 0, *σ* = 10 × 10^−7^ N m^−1^, *κ* = 400 × 10^−21^ J using different Δ values. Δ is defined as ratio of depth of field to the radius, for spherical vesicles; due to the cells’ biconcave shape, the vertical radius is half the height of the RBC, and Δ would be a large value (up to 0.5) in fluorescence. However, in brightfield imaging this is likely reduced due to the interaction of membrane geometry and the imaging, with more contrast being created when the membrane parallel to the optical axis, this being at the cell’s equator. The value of Δ = 0.15 is used in this work, as discussed in the text.

Following the standard definition for Full Width Half Maximum (FWHM) of the axial point spread function of a microscope (40, 41) gives:

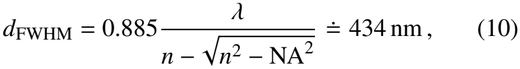

where we used the index of refraction of water and capped the effective NA of the objective at *n*_water_ = 1.33 due to the geometry at the interface. Rautu et al. (21) defined depth of field through standard deviation instead of FWHM, giving depth of field for the correction as *d* _*σ*_ = *d*_FWHM_ / 2.355 ≈ 184 nm. At a typical RBC height of 2.5 µm, this gives a Δ≈ 0.15.

Determining a precise value of Δ, linked to the NA of the objective, and all the optics including the condenser illumination, will require further study, and will potentially unlock the possibility of an absolute measurement of tension and bending modulus, consistent across different optical setups. What is clear from our work so far is that the focal depth effect is important in measuring RBC properties, and that since the measured values are sensitive to Δ, see Figure 5, the absolute values taken with different optical setups probably cannot be directly compared. For our setup, based on the calculation above, we use Δ = 0.15.

### 4.3 High-pass filtering attenuates non-thermal effects

The active effects on the membrane, introduced in Section 2.4, have been shown to have timescales around 100 ms (30, 35), with thermal behavior seen at frequencies above this (29, 35). The slowest modes used for fluctuation analysis in RBCs typically have decay times around 20 ms (22), enabling the possibility to isolate the two contributions.

To separate the two regimes, we apply a 4th-order high-pass Butterworth filter in both directions using SciPy (42) directly to the time series of the complex Fourier mode amplitudes *a*_*n*_ (*t*). The filtering is executed using a zero-phase forward-backward algorithm (`sosfiltfilt` in *SciPy*) to prevent any temporal phase shifts. Prior to applying the filter, the temporal mean of each mode (the time-independent static shape) is subtracted. This subtraction helps to prevent edge artifacts at the signal boundaries during filtering. The high-pass filter subsequently attenuates the potential contribution from active processes as well as any other slow changes, such as mechanical effects of flows or cell rotation. The effect of this filter on an example power spectrum can be seen in Figure 6: the entire spectrum is shifted lower as slow-timescale effects are suppressed. While static shape subtraction removes the perfectly constant component, the filter additionally eliminates the slow-timescale ‘bumps’ visible in the raw spectrum, which arise from slowly changing mean cell shape (e.g. rotation) or low-frequency active deformations that are not perfectly time-invariant.

**Figure 6:**
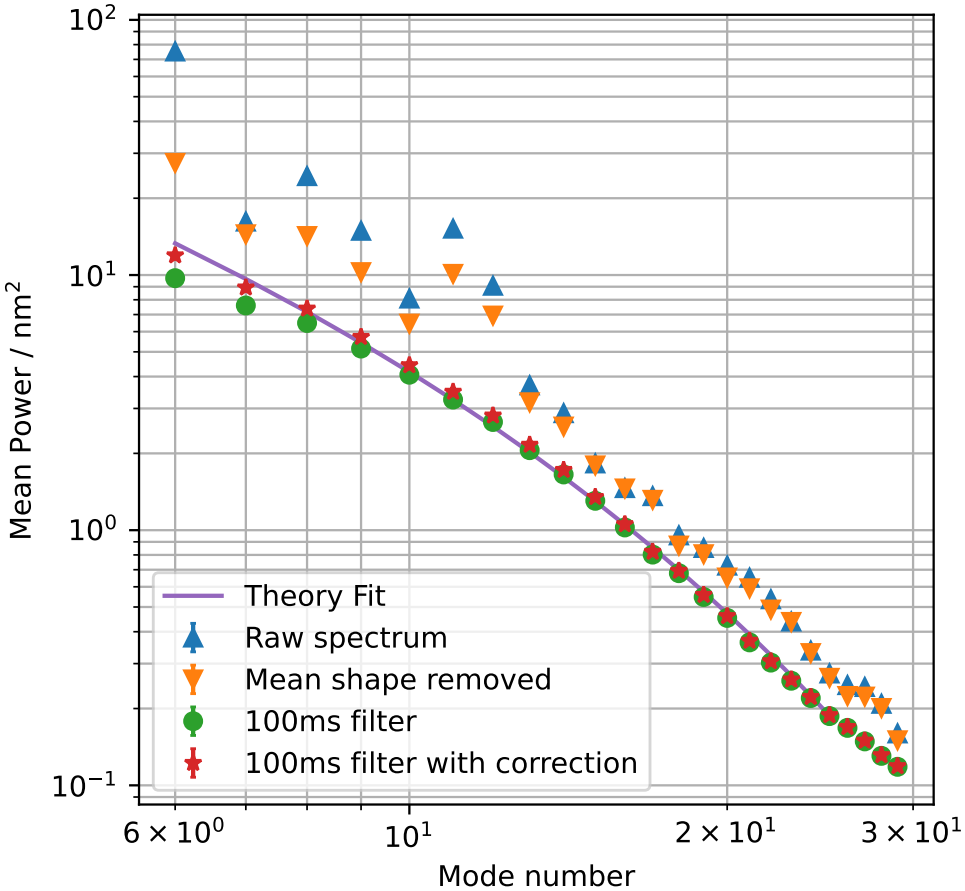
Effect of high pass filter and mean shape removal on spectrum obtained from a single cell in a commercial chamber in wash medium (Section 3.4). The static shape removal can help avoid the constant shape but will not remove slow changes. The filtered spectrum with the correction for power lost by filtering (as described in Section 4.3.1) matches the expected shape of the spectrum as seen in the fit using Eq. 4. In this figure the spectra are not corrected for depth of field or for exposure time - these corrections are included in the function used for the theory fit line.

#### 4.3.1 Accounting for the thermal fluctuation power loss due to filtering

Thermal modes with an exponential decay have a Lorentzian power spectrum which is maximal at zero frequency. A high-pass filter inevitably attenuates this spectrum by removing its central portion, making the measured power spectrum systematically lower than the ‘true’ power spectrum. This attenuation is shown in frequency space in Figure 7, and in the form of recovered/true spectrum on simulated data in Figure 8.

**Figure 7:**
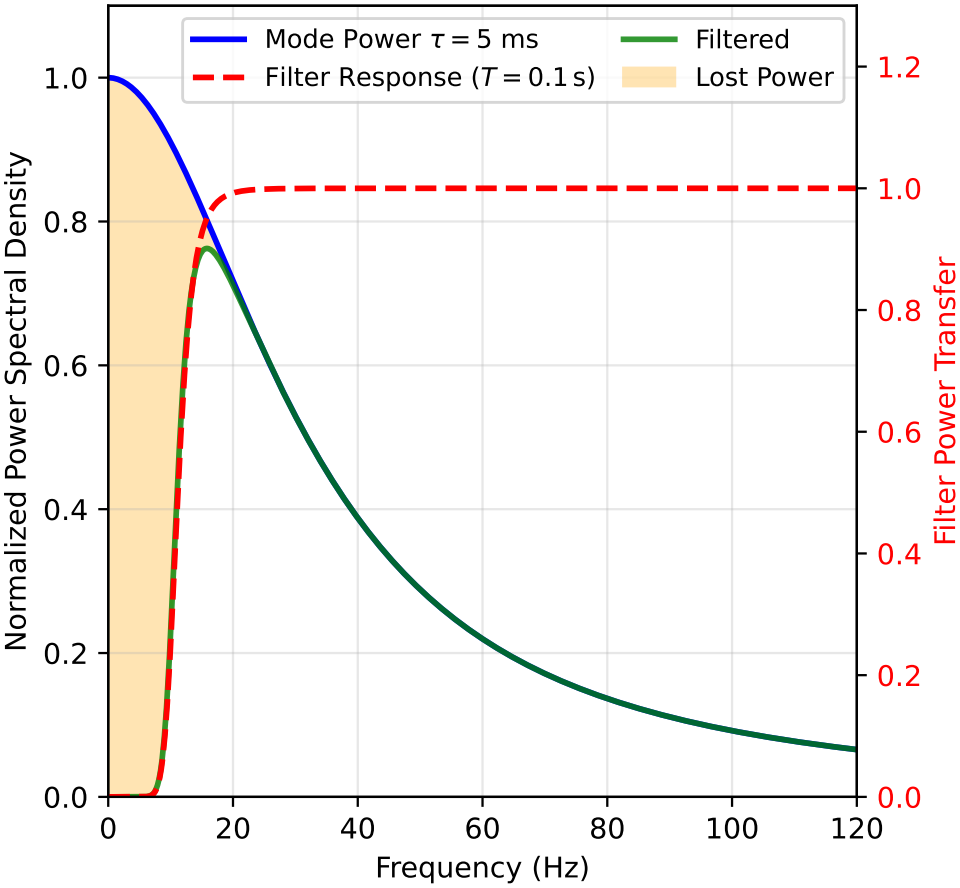
Effect of the high-pass filter in frequency space shown on a thermal mode with 5 ms decay time. The lost power in the low frequency component of the spectrum is shown highlighted on the left. Due to the shape of the spectrum this loss is unavoidable as in the case of a mixed active and thermal signal, this part of the spectrum mixes the two.

**Figure 8:**
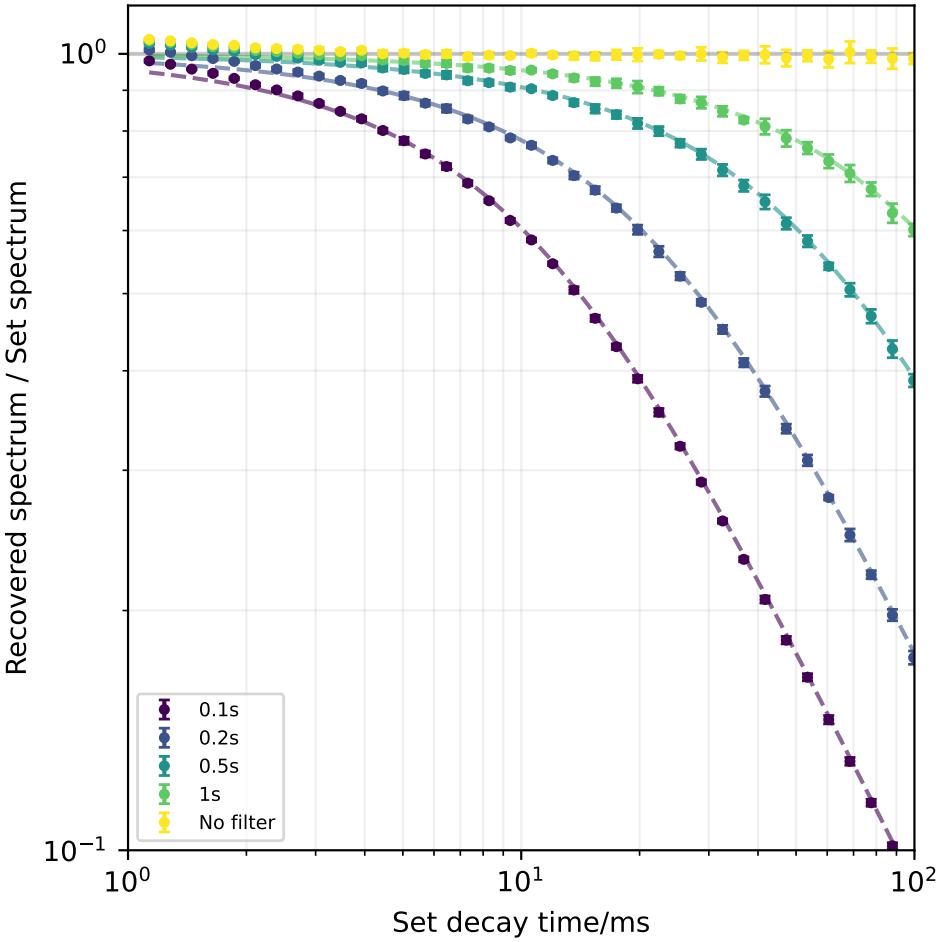
Power spectrum attenuation as a result of applying a high-pass Butterworth filter with different cut off times on simulated exponentially decaying excitations with different decay times. Dashed lines represent predicted attenuation based on filtering an exponentially decaying function. Inverse of this prediction is used to correct for the attenuation.

To recover the original power, a correction factor *K* must be applied to multiply the measured power of each mode. This correction factor is defined as the ratio of the total power of the unfiltered Lorentzian spectrum to the power remaining after zero-phase forward-backward Butterworth filtering of order *N* = 4. This is obtained by directly integrating the filter response over the Lorentzian spectrum of a mode with decay time *τ*, and depends only on the ratio of *τ* to the filter cutoff period *T*:

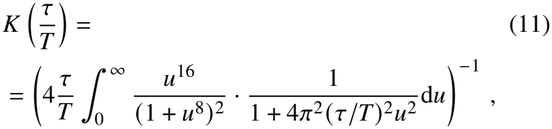

where *u* = *ω* / *ω*_*c*_ is the frequency normalized by the filter’s cutoff frequency *ω*_*c*_ = 2*π* / *T*. This integral is evaluated numerically for each mode’s decay time *τ* and applied to the power spectrum before fitting.

#### 4.3.2 Simulated active “kicks” to the membrane

To verify that the filter can remove the non-thermal effects we used the custom simulation introduced in Section 3.3, with the non-thermal mode generation enabled and amplitude of non-thermal modes set to match the difference between active and passive cells reported by Betz et al. (29) (48 %). This non-thermal contribution was largely eliminated from the fast modes when a high-pass filter with a cutoff matching their lifetime or faster was used as shown in Figure 9.

**Figure 9:**
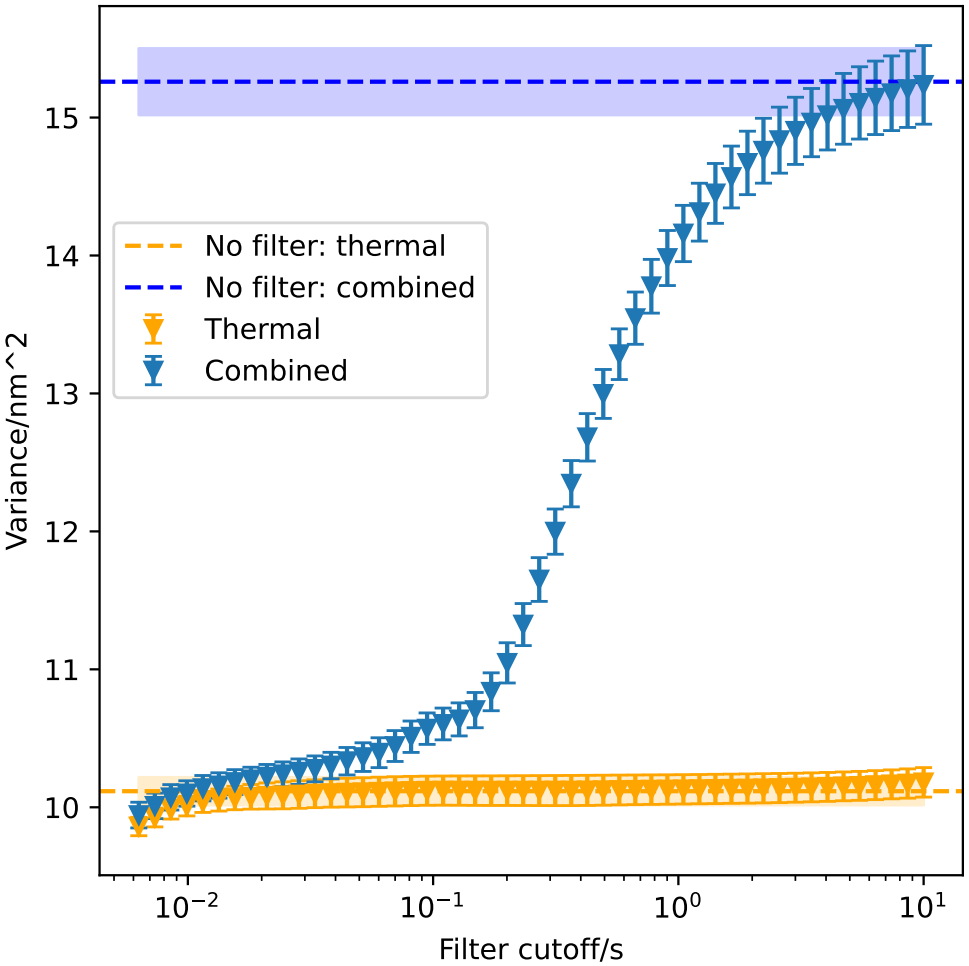
The effect of a high pass filter on simulated active contributions to flickering with a lifetime of *τ* = 100 ms. The filter is applied on the contour data (*r* (*ϕ, t*)) obtained from the simulation to test the entire pipeline. The decay times used for the correction for lost power are obtained from fits in the correlations as they would be when run with real data. The high pass filter can eliminate approximately 90 % of the non-thermal contribution from a mixed signal when the mode decay time is approximately equal to the filter cutoff.

The choice of a filter cutoff presents a trade-off between effective elimination of non-thermal modes and increased uncertainty due to increased reliance on the correction described in Section 4.3.1, which is itself limited by the error in obtained decay times of thermal modes. The minimum practical filter cutoff is also limited by the framerate used in experiments: recording at 660 fps we find that 100 ms presents a good tradeoff between these effects.

### 4.4 Accounting for mode dynamics with non-thermal contributions: Linear + exponential decay

The thermal modes are overdamped excitations (15). This results in an exponentially decaying autocorrelation of amplitudes of individual spherical modes, with the decay constant given by the decay time of the mode. This is the standard way of obtaining mode decay times and with some attention to the spherical-to-equatorial projection has also been used to obtain intracellular viscosity, for example in (22, 43).

The afore discussed non-thermal contributions and other slow changes can also be seen as a long tail in the autocorrelations. Under the assumption that the non-thermal contributions consist of ‘bumps’ to the membrane shape, which remain constant for a fixed lifetime before decaying as the corresponding thermal mode, this would contribute an additional linear term to the autocorrelation, which is consistent with our data, see Figure 10.

**Figure 10:**
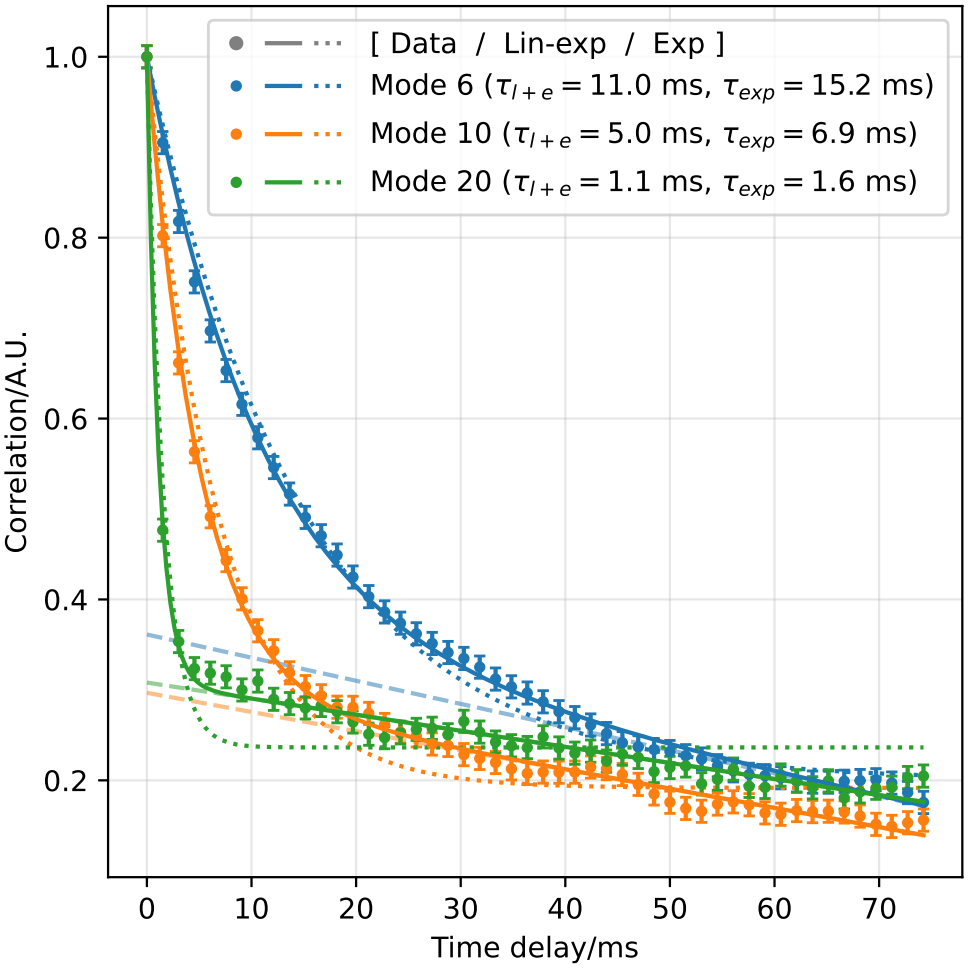
Autocorrelations are fitted well with the linear component included, compared to the traditional single exponential decay (dotted line). This data is from a single cell, imaged in a commercial chamber, in the standard wash medium. Dashed line shows the linear component. Using the linear + exponential model (solid lines) we can obtain better fits especially at higher modes, extending the range of modes usable for studying the dynamics. The modes shown are selected for display purposes; equivalent plots for all modes (6–20) are shown in Figure S7.

To validate the addition of the linear term when extracting timescales from autocorrelation functions we used our automated imaging system to record cells in a commercial chamber in wash medium, using different exposure times between 0.9 ms and 9.9 ms, and fitted the observed mode power attenuation using Eq. 2 (20). This is shown in Figure 11 for 3 modes from one cell with different decay times. This process provides a second source of mode decay times, effectively probing a much shorter timescale. By applying this method to a dataset of 863 cells, we obtained 17896 modes where both this method and the autocorrelation fitting yielded a decay time, allowing a comparison between these and showing that the linear + exponential model better matches results based on the camera shutter time (Figure 12). The simple exponential model tends to overestimate the thermal decay times due to the long tail.

**Figure 11:**
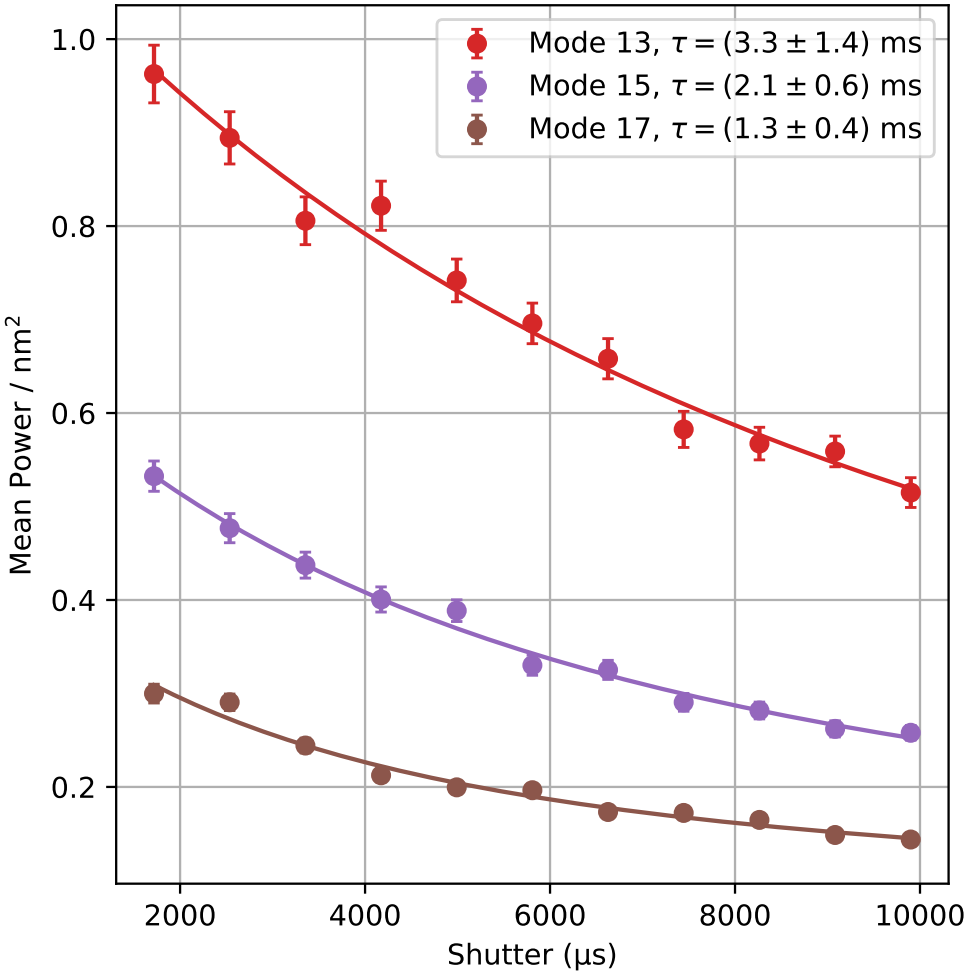
Power observed in 3 modes of a single cell at different exposure times shows the expected attenuation due to in-camera averaging. Fits using Eq. 2 determine the decay time of each mode. The modes shown are selected for display purposes; equivalent plots for all modes (6–20) are shown in Figure S8.

**Figure 12:**
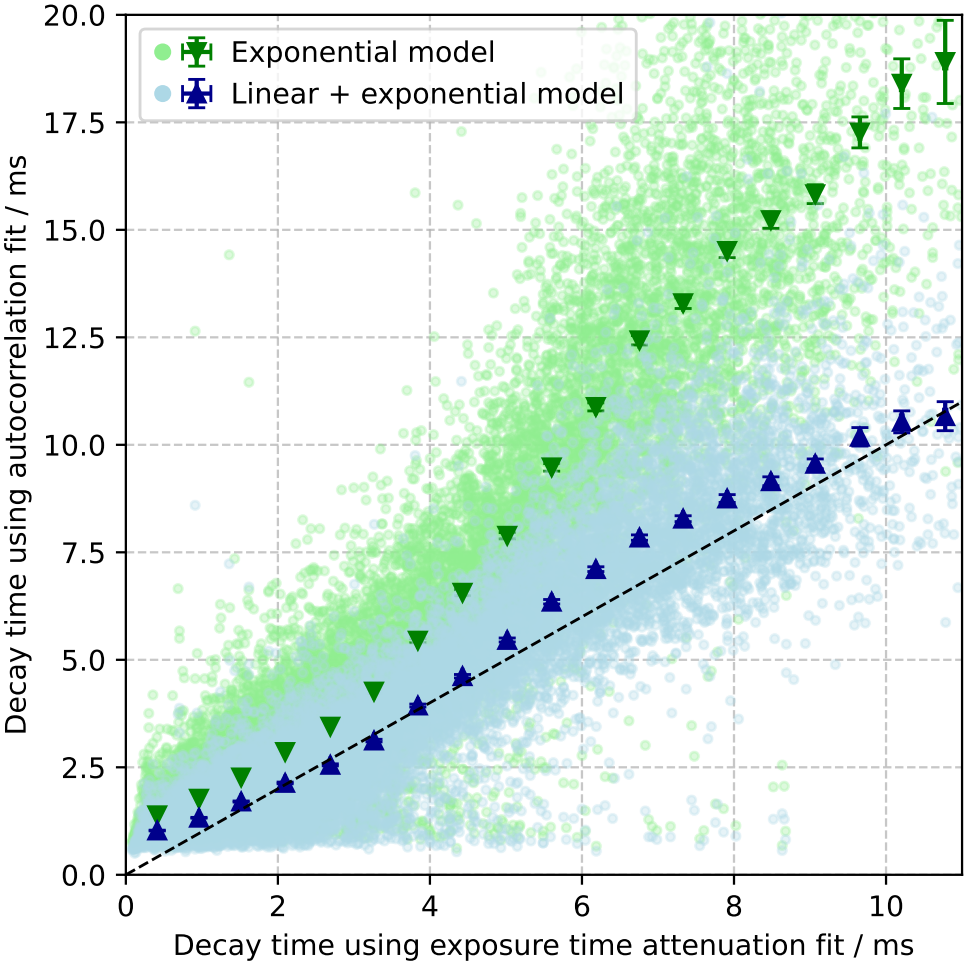
Comparing the two autocorrelation based results shows that the linear + exponential decay model (blue) results in decay times that agree well with those from the exposure time attenuation fits, whereas the more traditional simple exponential model (green) systematically overestimates the decay times. This comparison is based on decay times *τ*_*q*_ obtained using the two different approaches, on a total of 17 896 modes across 863 cells. The same cells were recorded at 12 different camera exposure times between 0.9 ms and 9.9 ms, with illumination power adjusted to maintain similar average intensity. In the first approach, the decay times are determined by fitting the attenuation of the observed mode power as a function of exposure time *T*_exp_ using the exposure time attenuation expression Eq. 2, example fits are shown in Figure 11. In the second approach, the decay times are extracted by fitting the temporal autocorrelation function of the mode amplitudes from a single high-speed video recorded at 660 fps (*T*_exp_ = 0.9 ms). We fitted the autocorrelations with an exponential and a linear + exponential decay model (Eq. 12), example fits are shown in Figure 10. Light points show individual modes, markers show the mean ± SEM within 25 equal-width bins of the exposure time fit decay time. The dashed black line shows equal decay times from the two approaches.

### 4.5 Fit range choice now has less effect

The pipeline described up to here significantly improves the issue of mode-range choice. Using the filtering together with the linear + exponential model of autocorrelations, and iteratively refining the fit of viscosity, tension, and bending modulus, both the spherical and planar fits become significantly less dependent on the choice of minimum mode included in the fit, as shown in Figure 13. At very low modes there is still some dependence in the planar case which is expected due to the planar approximation not being appropriate here. The spherical model, especially at 0 depth of field performs very well here down to mode 3.

**Figure 13:**
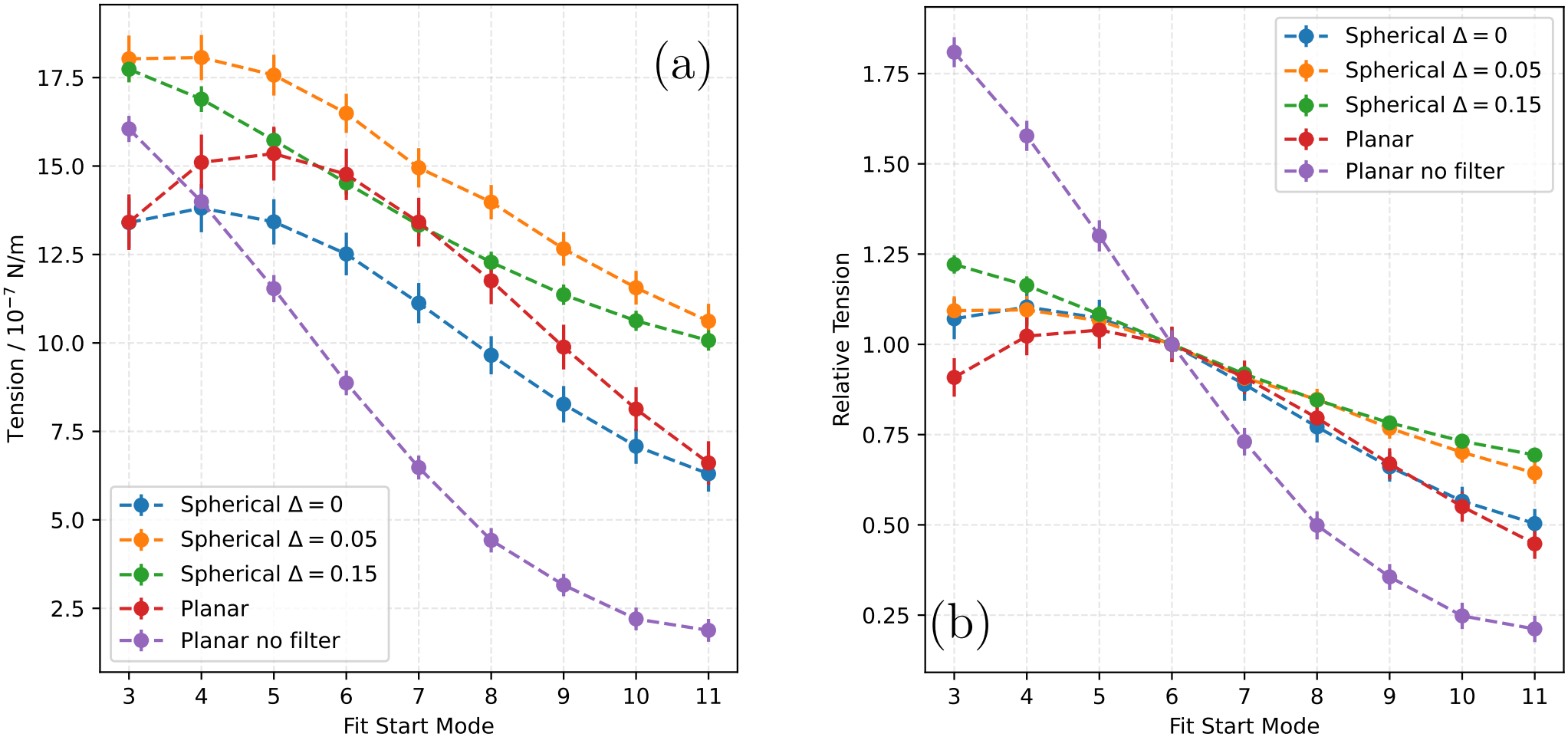
Effect of the choice of minimum mode to include in the fit on the obtained RBC tension for different configurations. Error bars show standard error in the mean across the 80 cells used in this comparison. Planar case involves fitting following Eq. 1 (20), the spherical fits use fits based on Eq. 4 (21). Both use camera exposure time compensation Eq. 2 (20) and the linear+exponential autocorrelation fits for decay times, fitted with the viscosity dependent expression from (22) and using the fitted results for shutter and filter effect compensation (see Section 4.7). Planar fitting uses a fixed maximum mode of 18 in the fit, spherical fits use automatic detection of when noise pushes the data above fit from lower modes per Section 4.7. 100 ms filter was applied to all but the last result - showing the filtering does make the result far more stable across a range of modes for both the spherical and planar fitting options. For this comparison the iterative refinement was run with damped decay time updates (Section S1.2.8 of the supporting material), as fits starting at high modes exclude the tension dominated modes and the three standard iterations do not converge there. (a) shows the effect on mean tension over 80 cells, in (b) the results are normalized to 1 at start mode 6 for all results, to highlight the changes.

The automatic selection of the maximum mode, based on when noise causes a separation between prediction based on lower maximum mode and the data, allows the fitting to use as wide a range as possible, i.e. maximizes the data used in the fit. This is essential as to obtain both tension and bending modulus from the fit, the range of modes should cover both the tension dominated regime and the bending dominated regime - the position of this crossover changes with the parameters and thus using as wide a range of modes is especially important in cases of high tension.

### 4.6 Glutaraldehyde increases tension

Glutaraldehyde has been shown previously to increase RBC tension without affecting bending modulus (13, 44) and this observation can be used to validate the pipeline. As expected, RBCs incubated with increasing concentrations of glutaralde-hyde show a significant increase in tension (Figure 14a) with minimal effect on the bending modulus (Figure 14b). These results were obtained following the optimized pipeline described in Section 4.7. This pipeline also produces an estimate of internal viscosity, this is shown in Figure S6. The relative change observed matches that reported in (13) for most concentrations (at 0.001 % glutaraldehyde we observe a 69 % increase while (13) reported 55 %). The difference and the faster increase at high concentrations can be explained by different methods of glutaraldehyde treatments.

**Figure 14:**
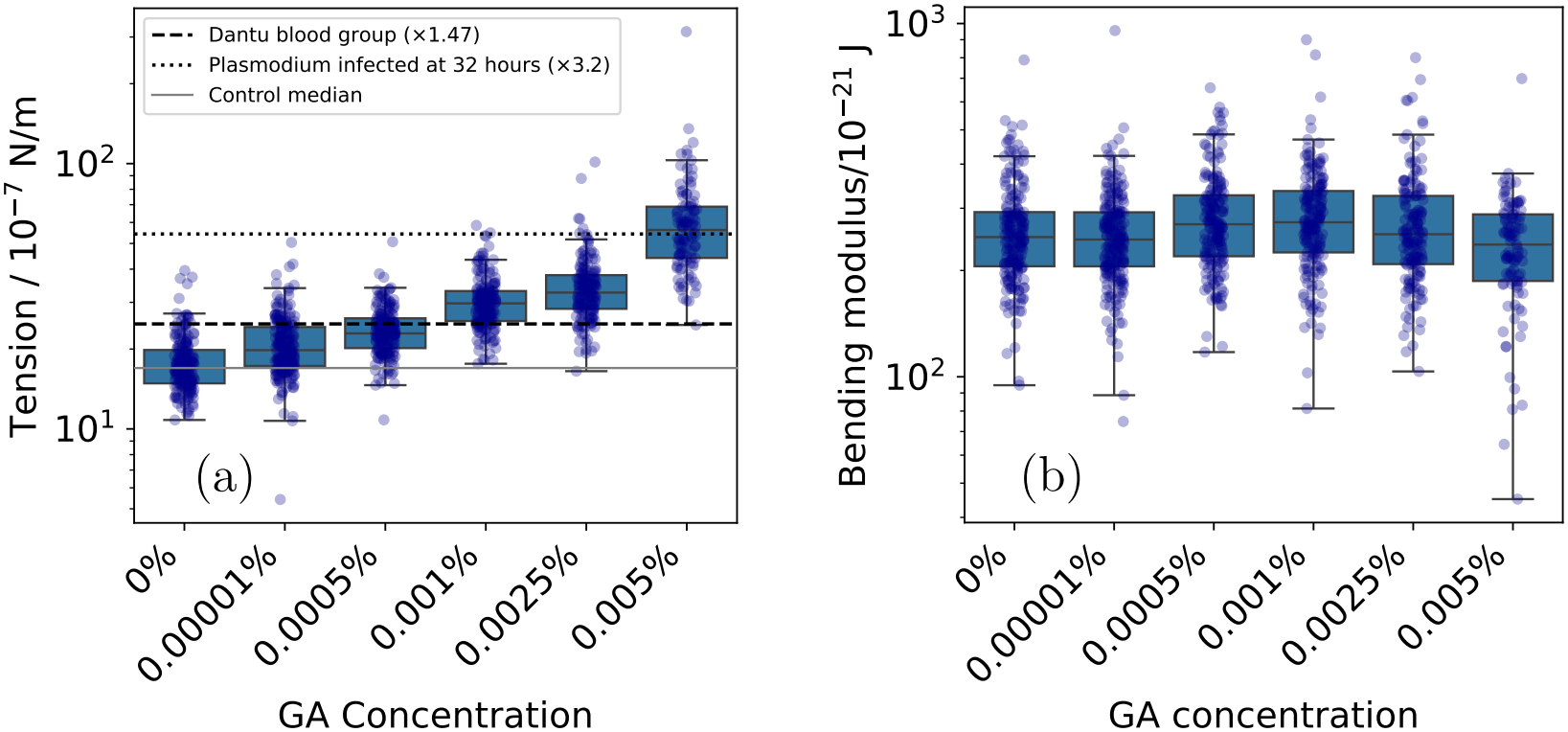
RBCs incubated for 30 min with different concentrations of glutaraldehyde show increase in tension (a) with no change in bending modulus (b), consistent with published results.(13) For comparison with biologically relevant changes, the horizontal lines in (a) show the control tension multiplied by the relative tension increase reported for Dantu homozygous RBCs (×1.47) (13) and for RBCs 32 h after infection with *Plasmodium falciparum* (×3.2) (38). The numbers of analyzed cells in each of the concentrations were 192, 213, 182, 176, 159, and 103 respectively.

These results are obtained by fitting the spherical fluctuation model with a non-zero depth of field correction (Δ = 0.15). The use of this correction appears to improve the separation between bending modulus and tension - without this correction we do see an concentration dependent increase in bending modules in addition to the tension increase. Result for this fit is in Figure S4.

### 4.7 Optimized fitting pipeline

Following the above discussed points, we have developed a self-consistent pipeline that combines the static shape fluctuations and dynamic decay properties of the membrane. The step-by-step pipeline is summarized below:

1. **Fourier Decomposition and Filtering :** The detected cell contour *r* (*ϕ, t*) is decomposed into 1D Fourier mode amplitude time series *u*_*q*_ (*t*). To remove slow active fluctuations and static shape features, a zero-phase high-pass Butterworth filter of order *N* = 4 with a cutoff period of *T* = 100 ms is applied to the time series *u*_*q*_ (*t*) (see Section 4.3.1). The mean power spectrum (MPS) is then calculated as the time-average of the squared magnitudes *⟨*|*u*_*q*_ |^2^⟩.
2. **Decay Time Extraction:** The autocorrelation functions (ACFs) of each mode amplitude *u*_*q*_ (*t*) are calculated and fitted using a linear-exponential model:

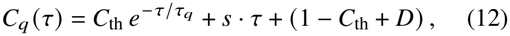

where *τ*_*q*_ is the fast thermal relaxation decay time (see Eq. S3 in the supporting material). This model successfully separates the thermal decay time *τ*_*q*_ from the slow active background.
3. **High-Pass Filter and Shutter Corrections:** The experimental MPS is corrected for two systematic attenuation effects:
  - Multiplied by the high-pass filter correction factor *K* (*τ*_*q*_ /*T*)(Eq. 11) to restore the power lost due to the high-pass filter.
  - Multiplied by the exposure time correction factor *C* (*τ*_*q*_ /*T*_exp_)(Eq. 2) to compensate for camera shutter temporal averaging.
  These initial corrections use the raw decay times *τ*_*q*_ from Step 2.
4. **Baseline Fitting and Auto-Cutoff Selection:** A baseline fit of the corrected MPS is performed over a narrow range of modes (*q∈* [6, 18]) using the projected spherical model (Eq. 4) to obtain initial estimates for tension *σ* and bending modulus *κ*. The maximum mode *q*_max_ for the full fit is determined automatically by finding where noise causes the measured MPS to systematically deviate above this baseline fit by more than 5% for at least 5 consecutive modes. The final fit is performed up to *q*_max_ − 1.
5. **Tension and Bending Modulus Fitting:** The corrected MPS is fitted over the range *q ∈* [6, *q*_max_ 1] using Eq. 4 to obtain tension *σ* and bending modulus *κ*, accounting for the depth-of-focus projection parameter Δ = 0.15.
6. **Viscosity and Decay Time Refinement:** The extracted raw decay times *τ*_*q*_ for modes whose timescales lie in the range 1.5 ms to 10 ms are fitted to the theoretical model (22):

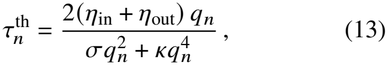

where *η*_out_ is the external medium viscosity, to extract the internal viscosity *η*_in_. During this fit, *σ* and *κ* are locked to the values obtained in Step 5, yielding a smoothed, noise-free set of fitted decay times 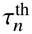 in addition to the viscosity estimate.
7. **Iterative refinement Loop:** To limit noise propagation from the raw decay times, the raw decay times *τ*_*q*_ in Step 3 are replaced by the fitted decay times 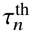 from Step 6. Steps 3 to 7 are then iterated 3 times to obtain the final parameters *σ, κ*, and *η*_in_.

Full details of the optimized analysis methodology are described in Section S1 of the supporting material.

## 5 CONCLUSION

In this work we have developed an improved method for using flickering spectroscopy on red blood cells. By applying appropriate high pass filtering, it is now possible to minimise the impact of non-thermal fluctuations. Using a linear + exponential model for autocorrelations, we can account for non-thermal contributions in the measurement of mode decay times. We have verified the improved performance of this method by observing perceived attenuation of fluctuations due to exposure time of the camera. Using a spherical fit together with exposure time compensation, filtering, and the linear-exponential decay time model we have greatly reduced the effect of the choice of which modes should be included in the fit - showing that this model better captures observed spectra.

We have verified the complete pipeline by recapitulating the tension increasing effect of glutaraldehyde.

In this pipeline we use a depth of field factor of Δ = 0.15, based on the depth of field of the optics used, and the typical height of an RBC. A direct measurement of this value in brightfield imaging is non-trivial. We established that this value can have a significant effect on the fit parameters and future work should focus on establishing a way of measuring this value directly for any imaging system used for flickering imaging experiments to ensure values obtained through different setups can be more directly compared.

Using this improved analysis method, together with our high-throughput automated imaging setup, flickering spectroscopy can now be used at scale to screen for compounds with potential effect on the membrane.

## DATA AND CODE AVAILABILITY

Analysis codes for the flickering pipeline are maintained on https://github.com/Cicuta-Group/flickering/ The version used for this work, and data for the figures in the manuscript, are on the Zenodo archive: https://doi.org/10.5281/zenodo.24367408

## AUTHOR CONTRIBUTIONS

F.A., J.C.R and P.C. contributed to the design and interpretation of experiments. F.A. and J.K. designed the equipment, and carried out experiments and analysis. F.A. drafted the manuscript, J.C.R and P.C. contributed. J.C.R and P.C. conceived and supervised the study. All authors discussed the results and contributed to the final version of the paper.

## ACKNOWLEDGMENTS

This work has benefited from discussions with Viola Introini, Sebastian Krauss, and Rini Ghosh. Guilherme Nettesheim and Jamie Morley contributed to early versions of the fitting code. Microscope interaction code benefited from in-house libraries developed by Boyko Vodenicharski and Morten Kals. This work was supported by the Engineering and Physical Sciences Research Council Centre for Doctoral Training in Sensor Technologies for a Healthy and Sustainable Future [EP/S023046/1].

## SUPPORTING MATERIAL

An online supplement to this article can be found by visiting BJ Online at http://www.biophysj.org.

## REFERENCES

1. Rahman, Y. E., D. L. Elson, and E. A. Cerny, 1973. Studies on the Mechanism of Erythrocyte Aging and Destruction I. Separation of Rat Erythrocytes According to Age by Ficoll Gradient Centrifugation. Mechanisms of Ageing and Development 2:141–150.

2. Ganzoni, A. M., R. Oakes, and R. S. Hillman, 1971. Red Cell Aging in Vivo. J Clin Invest 50:1373–1378.

3. Diez-Silva, M., M. Dao, J. Han, C.-T. Lim, and S. Suresh, 2010. Shape and Biomechanical Characteristics of Human Red Blood Cells in Health and Disease. MRS Bull 35:382–388.

4. Da Costa, L., J. Galimand, O. Fenneteau, and N. Mohandas, 2013. Hereditary spherocytosis, elliptocytosis, and other red cell membrane disorders. Blood Reviews 27:167–178.

5. Agrawal, R., T. Smart, J. Nobre-Cardoso, C. Richards, R. Bhatnagar, A. Tufail, D. Shima, P. H. Jones, and C. Pavesio, 2016. Assessment of red blood cell deformability in type 2 diabetes mellitus and diabetic retinopathy by dual optical tweezers stretching technique. Sci Rep 6:15873.

6. Lopes, C. S., A. C. Pronto-Laborinho, V. A. Conceição, T. Freitas, G. L. Matias, M. Gromicho, N. C. Santos, M. de Carvalho, and F. A. Carvalho, 2024. Erythrocytes’ surface properties and stiffness predict survival and functional decline in ALS patients. BioFactors 50:558–571.

7. Strijkova-Kenderova, V., S. Todinova, T. Andreeva, D. Bogdanova, A. Langari, A. Danailova, S. Krumova, E. Zlatareva, N. Kalaydzhiev, I. Milanov, and S. G. Taneva, 2021. Morphometry and Stiffness of Red Blood Cells-Signatures of Neurodegenerative Diseases and Aging. International Journal of Molecular Sciences 23:227.

8. Gomi, T., T. Ikeda, and F. Ikegami, 1997. Beneficial Effect of α-Blocker on Hemorheology in Patients With Essential Hypertension. Am J Hypertension 10:886–892.

9. Lau, C. S., A. R. Saniabadi, and J. J. F. Belch, 1995. Reduced red blood cell deformability in patients with rheumatoid vasculitis improvement after in vitro treatment with dipyridamole. Arthritis & Rheumatism 38:248–253.

10. Saha, A. K., B. R. Schmidt, J. Wilhelmy, V. Nguyen, A Abugherir, J. K. Do, M. Nemat-Gorgani, R. W. Davis, and A. K. Ramasubramanian, 2019. Red blood cell de-formability is diminished in patients with Chronic Fatigue Syndrome. Clinical Hemorheology and Microcirculation 71:113–116.

11. Tsukada, K., E. Sekizuka, C. Oshio, and H. Minamitani, 2001. Direct Measurement of Erythrocyte Deformability in Diabetes Mellitus with a Transparent Microchannel Capillary Model and High-Speed Video Camera System. Microvascular Research 61:231–239.

12. Moon, J. S., J. H. Kim, J. H. Kim, I. R. Park, J. H. Lee, H. J. Kim, J. Lee, Y. K. Kim, J. S. Yoon, K. C. Won, and H. W. Lee, 2016. Impaired RBC deformability is associated with diabetic retinopathy in patients with type 2 diabetes. Diabetes & Metabolism 42:448–452.

13. Kariuki, S. N., A. Marin-Menendez, V. Introini, B. J. Ravenhill, Y.-C. Lin, A. Macharia, J. Makale, M. Tendwa, W. Nyamu, J. Kotar, M. Carrasquilla, J. A. Rowe, K. Rockett, D. Kwiatkowski, M. P. Weekes, P. Cicuta, T. N. Williams, and J. C. Rayner, 2020. Red Blood Cell Tension Protects against Severe Malaria in the Dantu Blood Group. Nature 585:579–583.

14. Browicz, V. T., 1890. Weitere Beobachtungen über Bewegungsphänomene an roten Blutkörperchen in pathologischen Zustände. Verlag, Berlin.

15. Brochard, F., and J. F. Lennon, 1975. Frequency Spectrum of the Flicker Phenomenon in Erythrocytes. J de Physique 36:1035–1047.

16. Helfrich, W., and R. M. Servuss, 1984. Undulations, Steric Interaction and Cohesion of Fluid Membranes. Il Nuovo Cimento D 3:137–151.

17. Burton, A. L., W. L. Anderson, and R. V. Andrews, 1968. Quantitative Studies on the Flicker Phenomenon in the Erythrocytes. Blood 32:819–822.

18. Fricke, K., and E. Sackmann, 1984. Variation of Frequency Spectrum of the Erythrocyte Flickering Caused by Aging, Osmolarity, Temperature and Pathological Changes. Biochimica Et Biophysica Acta 803:145–152.

19. Zilker, A., M. Ziegler, and E. Sackmann, 1992. Spectral Analysis of Erythrocyte Flickering in the 0.3–4 µm Regime by Microinterferometry Combined with Fast Image Processing. Phys. Rev. A 46:7998–8001.

20. Pécréaux, J., H.-G. Döbereiner, J. Prost, J.-F. Joanny, and P. Bassereau, 2004. Refined Contour Analysis of Giant Unilamellar Vesicles. Eur. Phys. J. E 13:277–290.

21. Rautu, S. A., D. Orsi, L. Di Michele, G. Rowlands, P. Cicuta, and M. S. Turner, 2017. The Role of Optical Projection in the Analysis of Membrane Fluctuations. Soft Matter 13:3480–3483.

22. Yoon, Y.-Z., H. Hong, A. Brown, D. C. Kim, D. J. Kang, V. L. Lew, and P. Cicuta, 2009. Flickering Analysis of Erythrocyte Mechanical Properties: Dependence on Oxygenation Level, Cell Shape, and Hydration Level. Biophys J 97:1606–1615.

23. Koch, M., K. E. Wright, O. Otto, M. Herbig, N. D. Salinas, N. H. Tolia, T. J. Satchwell, J. Guck, N. J. Brooks, and J. Baum, 2017. Plasmodium Falciparum Erythrocyte-Binding Antigen 175 Triggers a Biophysical Change in the Red Blood Cell That Facilitates Invasion. Proc Natl Acad Sci USA 114:4225–4230.

24. Evans, J., W. Gratzer, N. Mohandas, K. Parker, and J. Sleep, 2008. Fluctuations of the Red Blood Cell Membrane: Relation to Mechanical Properties and Lack of ATP Dependence. Biophys. J. 94:4134–4144.

25. Helfrich, W., 1973. Elastic Properties of Lipid Bilayers: Theory and Possible Experiments. Zeitschrift für Naturforschung C 28:693–703.

26. Parpart, A. K., and J. H. F. Hoffman, 1956. Flicker in Erythrocytes. “Vibratory Movements in the Cytoplasm”? J Cell Comparative Physiol 47:295–303.

27. Boss, D., A. Hoffmann, B. Rappaz, C. Depeursinge, P. J. Magistretti, D. V. de Ville, and P. Marquet, 2012. Spatially-Resolved Eigenmode Decomposition of Red Blood Cells Membrane Fluctuations Questions the Role of ATP in Flickering. PLOS ONE 7:e40667.

28. Szekely, D., T. W. Yau, and P. W. Kuchel, 2009. Human Erythrocyte Flickering: Temperature, ATP Concentration, Water Transport, and Cell Aging, plus a Computer Simulation. Eur Biophys J 38:923–939.

29. Betz, T., M. Lenz, J.-F. Joanny, and C. Sykes, 2009. ATP-dependent Mechanics of Red Blood Cells. Proc. Natl. Acad. Sci. USA 106:15320–15325.

30. Gov, N. S., and S. A. Safran, 2005. Red Blood Cell Membrane Fluctuations and Shape Controlled by ATP-Induced Cytoskeletal Defects. Biophys J 88:1859–1874.

31. Tuvia, S., A. Almagor, A. Bitler, S. Levin, R. Korenstein, and S. Yedgar, 1997. Cell Membrane Fluctuations Are Regulated by Medium Macroviscosity: Evidence for a Metabolic Driving Force. Proc Nat Acad Sci USA 94:5045–5049.

32. Caselli, N., M. García-Verdugo, M. Calero, N. Hernando-Ospina, J. A. Santiago, D. Herráez-Aguilar, and F. Monroy, 2024. Red Blood Cell Flickering Activity Locally Controlled by Holographic Optical Tweezers. iScience 27.

33. Hernando-Ospina, N., M. Calero, G. Solís, I. G. Azcárate, D. Herráez-Aguilar, N. Caselli, L. H. Moleiro, J.-C. Segovia, J. M. Bautista, and F. Monroy, 2026. Metabolic Rigidity as a Mechanical Barrier to Malaria: Flickering Loss in PKLR-Deficient Erythrocytes. FASEB J 40:0892–6638.

34. Hernando-Ospina, N., M. Calero, L. H. Moleiro, D. Herráez-Aguilar, and F. Monroy, 2026. Structured volatility reveals dissipative lifelines in red blood cells. Phys. Rev. E 114:054409.

35. Turlier, H., D. A. Fedosov, B. Audoly, T. Auth, N. S. Gov, C. Sykes, J.-F. Joanny, G. Gompper, and T. Betz, 2016. Equilibrium Physics Breakdown Reveals the Active Nature of Red Blood Cell Flickering. Nature Physics 12:513–519.

36. Rodríguez-García, R., I. López-Montero, M. Mell, G. Egea, N. S. Gov, and F. Monroy, 2015. Direct Cytoskeleton Forces Cause Membrane Softening in Red Blood Cells. Biophys J 108:2794–2806.

37. Knotz, G., T. M. Muenker, T. Betz, and M. Krüger, 2026. Time reversal breaking of colloidal particles in cells. J. Chem. Phys. 165:015102.

38. Davies, H., H. Belda, M. Broncel, X. Ye, C. Bisson, V. Introini, D. Dorin-Semblat, J.-P. Semblat, M. Tibúrcio, B. Gamain, M. Kaforou, and M. Treeck, 2020. An Exported Kinase Family Mediates Species-Specific Erythrocyte Remodelling and Virulence in Human Malaria. Nature Microbiol 5:848–863.

39. Kals, M., L. Mancini, J. Kotar, A. Donald, and P. Cicuta, 2024. Multipad Agarose Plate: A Rapid and High-Throughput Approach for Antibiotic Susceptibility Testing. J Roy Soc Interface 21:20230730.

40. Lalaoui, L., M. Bouafia, D. Issaad, and A. Medjahed, 2020. Axial Point Spread Function Modeling by Application of Effective Medium Approximations: Widefield Microscopy. Optik 212:164646.

41. Sheppard, C. J. R., 1988. Depth of field in optical microscopy. J Microscopy 149:73–75.

42. Jones, E., T. Oliphant, P. Peterson, et al., 2001–. SciPy: Open source scientific tools for Python. http://www.scipy.org/.

43. Popescu, G., T. Ikeda, K. Goda, C. A. Best-Popescu, M. Laposata, S. Manley, R. R. Dasari, K. Badizadegan, and M. S. Feld, 2006. Optical Measurement of Cell Membrane Tension. Phys Rev Lett 97:218101.

44. Baskurt, O. K., M. Hardeman, M. Uyuklu, P. Ulker, M. Cengiz, N. Nemeth, S. Shin, T. Alexy, and H. J. Meiselman, 2009. Comparison of Three Commercially Available Ektacytometers with Different Shearing Geometries. Biorheology 46:251–264.

