## Supporting materials for "Robust and High-Throughput Flickering Spectroscopy for Measurement of Red Blood Cell Membrane Mechanics"

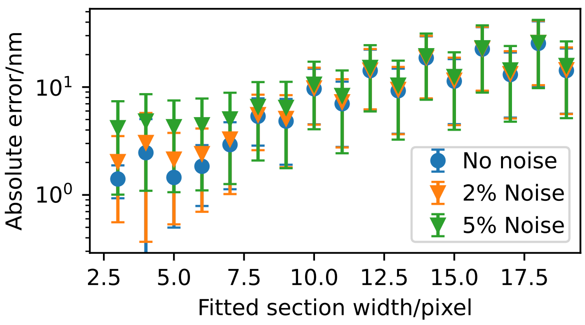


Figure S1: Recovery of artificially shifted profiles. As the shape of the correlation peak is not parabolic, precision decreases at large fit widths. Fit width of 5 was chosen as optimal as it provides some tolerance to random errors affecting a single point with no loss of precision.

### S1 DETAILS OF THE TRACKING AND FITTING PROCEDURE

#### S1.1 Contour Tracking

The tracking algorithm processes each video frame to extract the equatorial contour of a single, approximately circular cell.

##### S1.1.1 Cell Centre Detection

The cell centre is initially determined using a Hough Circle Transform applied to a denoised version of each image. The raw grayscale frame is normalised and denoised using a non-local means filter. The circle whose centre is closest to the image centre is selected as the initial estimate. To improve this estimate, a refinement step is performed: the full contour detection pipeline is executed temporarily with a fixed Gaussian-gradient based correlation mask and iterative centre re-estimation until the centre position converges within a tolerance of 0.005 pixels. This refined centre is then fixed as an initial estimate for all subsequent per-frame contour tracking. For the first image, this refined centre is used to generate the final correlation mask.

##### S1.1.2 Polar Image Unrolling

Each image is transformed from Cartesian to polar coordinates centred on the cell centre using bilinear interpolation. The polar image consists of 360 angular rays spanning the full $2\pi$ range and extends radially over a band of 60 pixels centred on the expected contour position. This ensures the membrane contour remains within the unrolled image even for slightly eccentric cells.

##### S1.1.3 Correlation Mask Generation

A one-dimensional correlation template (mask) is constructed from the angular-averaged radial intensity profile of the unrolled image. To suppress background bias and improve the discriminating power of the cross-correlation, the mean intensity of the non-mask regions is subtracted from both the template and the unrolled image. This mask is computed from the first frame and reused for all subsequent frames. To further improve mask quality, a refinement step is applied where the image is re-unrolled around the initially detected contour, and the resulting angular-averaged profile provides the final refined template. This step significantly increases the mask sharpness for less circular cells.

##### S1.1.4 Contour Detection via Sliding Cross-Correlation

The contour is located in the unrolled image by maximising the cross-correlation between the template and each radial column:

1. **Sliding correlation:** For each azimuthal angle, the dot product of the template with every possible radial window in the unrolled image is computed.

2. **Continuity constraint:** The correlation maximum at each angle is found within a restricted window around the running average of the previous angles. This prevents the contour from jumping to imaging artifacts or nearby cells.

3. **Sub-pixel refinement:** A second-order polynomial is fitted to a 5-pixel window around each correlation maximum. The vertex of the parabola provides the sub-pixel radial position for that angle.

##### S1.1.5 Refinement and Centre Convergence

After initial detection, a two-phase refinement is performed. The original image is re-unrolled using bicubic interpolation, centred on the detected contour. This places the membrane at the centre of the unrolled image for every angle, providing more uniform radial sampling. A second pass of the correlation-based detection is then performed. Finally, the cell centre is updated based on the mean position of the detected contour, and the process iterates until the centre shift converges within a tolerance of 0.05 pixels.

##### S1.1.6 Contour Validation

Each detected contour is subjected to quality checks based on the percentile range of radii, standard deviation, and the maximum Laplacian of the smoothed contour. Frames failing any of these criteria are marked invalid and excluded from subsequent spectral analysis.

#### S1.2 Contour Analysis and Fitting

The time series of radial contours is used to compute the fluctuation power spectrum and to extract decay times.

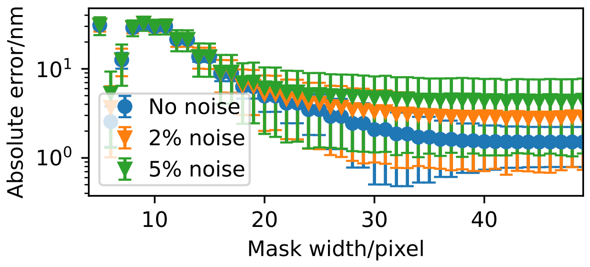


Figure S2: Effect of mask width on the precision, with added noise the benefits of a longer mask decrease and given typical noise values 30 pixels was chosen.

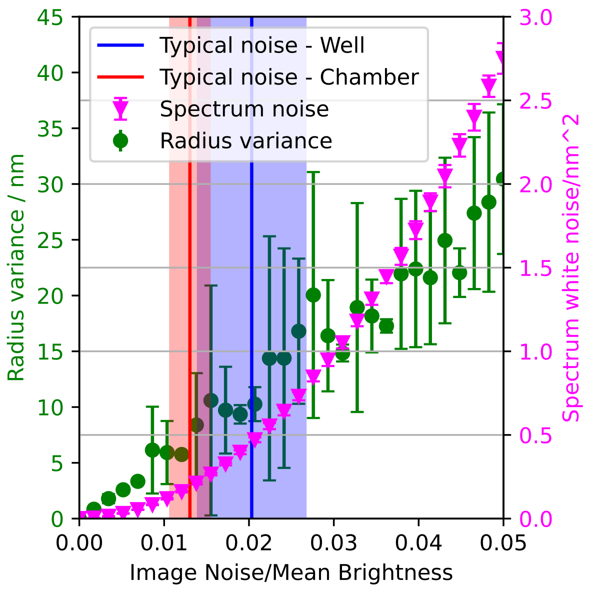


Figure S3: Tracking a fixed image with added noise. Videos recorded in a chamber normally have less noise as the wells of a well plate reduce illumination NA. Lower noise does reduce the spectrum noise floor, which enables the use of higher $q$ modes in the analysis.

##### S1.2.1 Fourier Decomposition

For each valid frame, the contour $r(\theta)$ is decomposed into angular Fourier modes:

$$\tilde{u}_q = \frac{1}{N} \sum_{j=0}^{N-1} r(\theta_j)\, e^{-2\pi i q j/N} \tag{S1}$$

where $N = 360$ and $q$ is the mode number. The mean radius is computed from the zeroth mode ($q = 0$).

##### S1.2.2 Static Shape Subtraction

To isolate thermal fluctuations from slow drifts and the static cell shape, a 4th-order Butterworth high-pass filter is applied to each Fourier mode time series. The filter has a cutoff period of 66 frames (100 ms). Zero-phase forward-backward filtering is used to avoid phase shifts. Prior to filtering, the temporal mean of each mode is subtracted.

##### S1.2.3 Mean Power Spectrum

The mean power spectrum (MPS) is computed as the time-average of the squared magnitudes of the Fourier coefficients:

$$\langle |u_q|^2 \rangle = \frac{1}{M} \sum_{t=1}^{M} |\tilde{u}_q(t)|^2 \tag{S2}$$

where $M$ is the number of valid frames. The uncertainty on each mode is estimated as the standard error of the mean.

##### S1.2.4 Decay Time Estimation

The temporal dynamics of each mode are characterised by fitting the autocorrelation function (ACF). The signal is mean-subtracted and normalised such that $C_q(0) = 1$. The ACF is fitted with a linear-exponential model:

$$C_q(\tau) = C_{\text{th}}\, e^{-\alpha\tau} + s \cdot \tau + (1 - C_{\text{th}} + D) \tag{S3}$$

where $\alpha = 1/\tau_q$ is the inverse decay time, $C_{\text{th}}$ is the thermal amplitude, $s$ is a linear drift term, and $D$ is a constant offset. The decay time $\tau_q$ is extracted for each mode.

##### S1.2.5 Viscosity-Fitted Decay Times

The experimental decay times are fitted to a theoretical model to extract the internal viscosity $\eta_{\text{in}}$ and to produce smoothed decay times for corrections. The model follows the expression derived by Yoon et al. (1):

$$\tau_n^{\text{th}} = \frac{2(\eta_{\text{in}} + \eta_{\text{out}})\, q}{\sigma q^2 + \kappa q^4} \tag{S4}$$

where $q = n/R$ is the wavevector and $\eta_{\text{out}}$ is the external medium viscosity. The tension $\sigma$ and bending modulus $\kappa$ are held fixed to the values obtained from the power spectrum fit.

##### S1.2.6 Spectrum Corrections

Two primary corrections are applied to the MPS:

1. **High-pass filter correction:** Compensates for the suppression of power by the high-pass filter. The correction factor is calculated as the ratio of total power to passed power for a Lorentzian spectral density filtered by the Butterworth response. In code this is referred to as rolling window correction.

2. **Exposure time correction:** Accounts for temporal averaging during the finite camera exposure ($T_{\text{exp}}$). Following Pécréaux et al. (2), the correction factor is $C(t) = 2t^2(1/t + e^{-1/t} - 1)$ where $t = \tau_q/T_{\text{exp}}$.

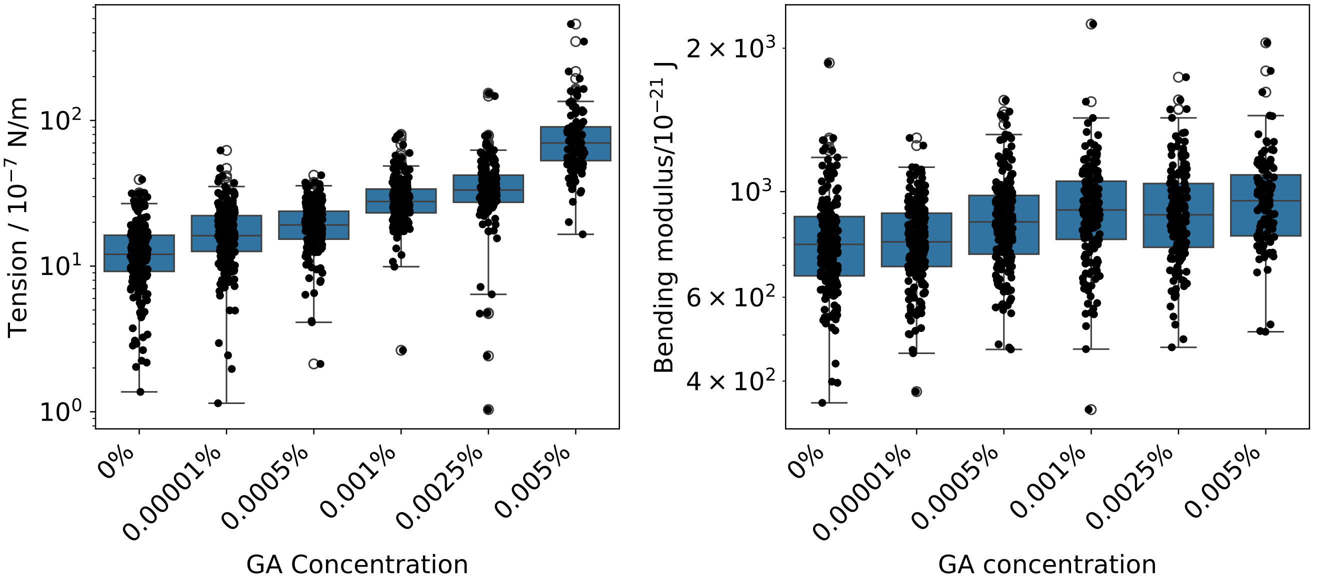


Figure S4: RBCs incubated for 30 min with different concentrations of glutaraldehyde and analyzed without a depth of field correction show increase in tension but an increase in bending modulus is also observed.

##### S1.2.7 Power Spectrum Fitting

The corrected MPS is fitted to the theoretical model developed by Rautu et al. (3), which accounts for depth-of-focus projection and assumes spherical harmonic structure of the fluctuations

$$\langle |u_q|^2 \rangle = 4\pi\alpha\beta^3 \sum_{n=q}^{q+30} \frac{|Y_q^n(0,\pi/2)|^2 L_{n,q}^2(\Delta) \cdot A_n}{(n-1)(n+2)[\beta^2 + n(n+1)]}, \quad \text{(S5)}$$

where $\alpha = k_B T/(4\pi\sigma\beta)$ and $\beta = R\sqrt{\sigma/\kappa}$. The factor $L_{n,q}(\Delta)$ accounts for the depth-of-focus projection.

The maximum mode $q_{\max}$ for the fit is determined automatically. A baseline fit is performed on a narrow range of low modes ($q \in [6, 18]$), and residuals are monitored for systematic deviations exceeding 5% for 5 consecutive modes, which defines the fit cutoff. The final fit is performed up to the mode one before where this deviation starts.

##### S1.2.8 Iterative Refinement

The fitting procedure iterates up to three times. After each fit, the decay times are re-estimated using the updated $\sigma$ and $\kappa$ values, and the spectral corrections are re-applied.

For the comparison of fit ranges (Figure 13 of the main text) this iteration was run 20 times with damped decay time updates: the decay times used in iteration $i+1$ are $\tau^{(i+1)} = (\tau^{\text{th}})^{0.3}(\tau^{(i)})^{0.7}$, where $\tau^{\text{th}}$ are the fitted decay times of iteration $i$. When the fit starts above mode 8 the spectrum no longer constrains the tension well and the undamped iteration can alternate between two solutions; the damped iteration settles on the self-consistent one. At the standard start mode of 6 both give the same mean tension within 2 %.

#### S1.3 Experimental Parameters

To match our imaging conditions we use the following parameters as standard: a frame rate of 660 fps, camera exposure time of 0.8 ms, and a pixel size of 98.6 nm/px. The temperature is set to 37 C, and the external viscosity is $0.733 \times 10^{-3}$ Pa s.

### S2 CONTOUR TRACKING PRECISION

To optimize the parameters we generated radial profile with artificial offset and tested how precisely the system can recover set offset. The procedure was

- Obtain radial intensity profile obtained from a real cell
- Interpolate this profile
- Apply a radial offset to this profile (i.e. shift it in the radial direction)
- Resample this profile based on the original resolution
- Apply the correlation based detection on this shifted profile using the original mask.

We used different mask widths and size of the area used for parabolic fit, with results shown in figures S1 and S2. The optimal choice appears to be fit width of 5 pixels (2 pixels each side of maximum) and performance improves with mask width, with 30 pixels chosen.

To determine the precision of this method on full 2D images, static images with added noise were tracked, with 10 frames from 16 cells each used to generate 1000 images by adding normally distributed random noise in different amounts. The results, shown in Figure S3, show the variance in the static contour to be around 7 nm for typical noise levels in our videos in the chambers, and that the expected errors do increase significantly with noise.

#### S3 ZERO-DEPTH OF FIELD GLUTARALDEHYDE ANALYSIS

When analyzing the results of the glutaraldehyde experiment with depth of field correction disabled (equivalent to the zero depth-of-field model of Pécréaux et al. (2)), a concentration-dependent increase in the resulting bending modulus is observed. This is contrary to published results (4) and the physical expectation that glutaraldehyde treatment only increases membrane tension without affecting the bending modulus.

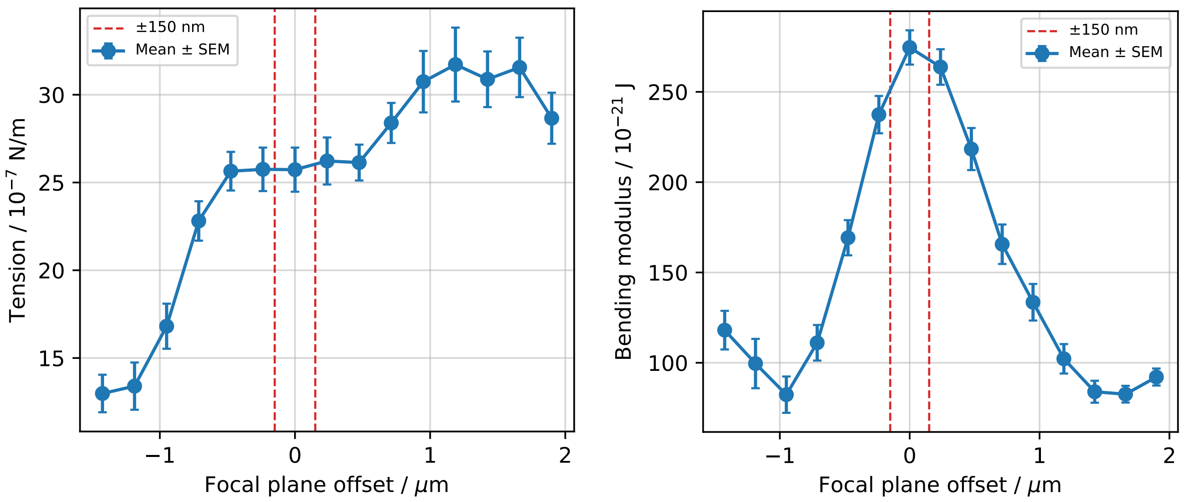


Figure S5: Effect of the focal plane position on obtained fit parameters. With an autofocus step size of 300 nm the image closest to the equator is within 150 nm, giving the worst-case error due to defocus of ≈ 8.6 % in bending modulus. The parabolic fit applied on the focus metrics to obtain a position in between the steps significantly reduces this error.

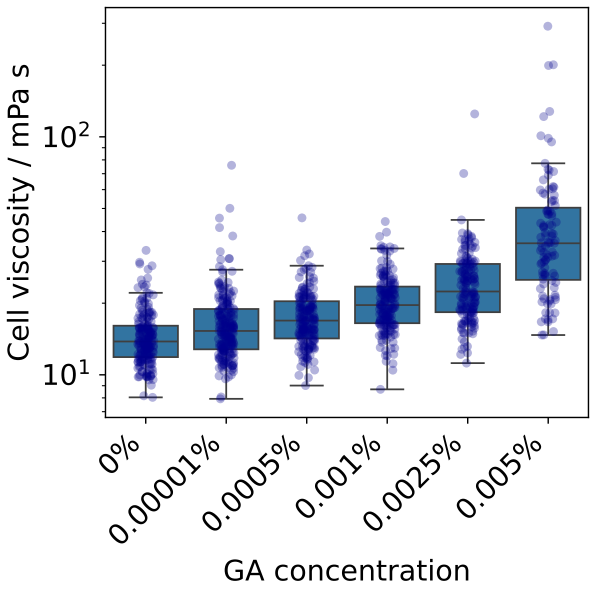


Figure S6: RBCs incubated for 30 min with different concentrations of glutaraldehyde show an increase in viscosity. The viscosity reported here is obtained by fitting of mode decay times using Eq. 13

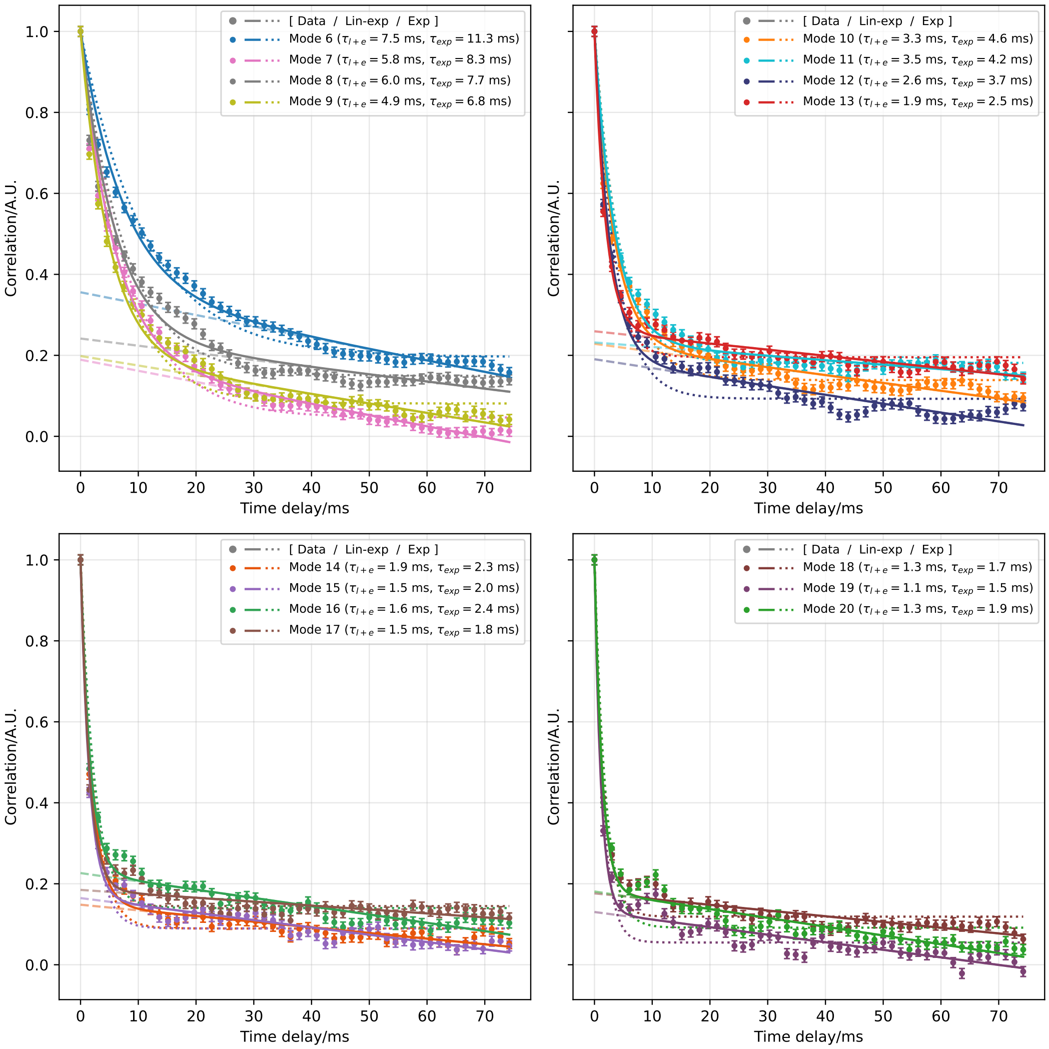


Figure S7: Autocorrelations are fitted well with the linear component included, compared to the traditional single exponential decay (dotted line). This data is from a single cell, imaged in a commercial chamber, in the standard wash medium. Dashed line shows the linear component. Using the linear + exponential model (solid lines) we can obtain better fits especially at higher modes, extending the range of modes usable for studying the dynamics. These are fits for a wide range of modes for the same cell as Figure 10

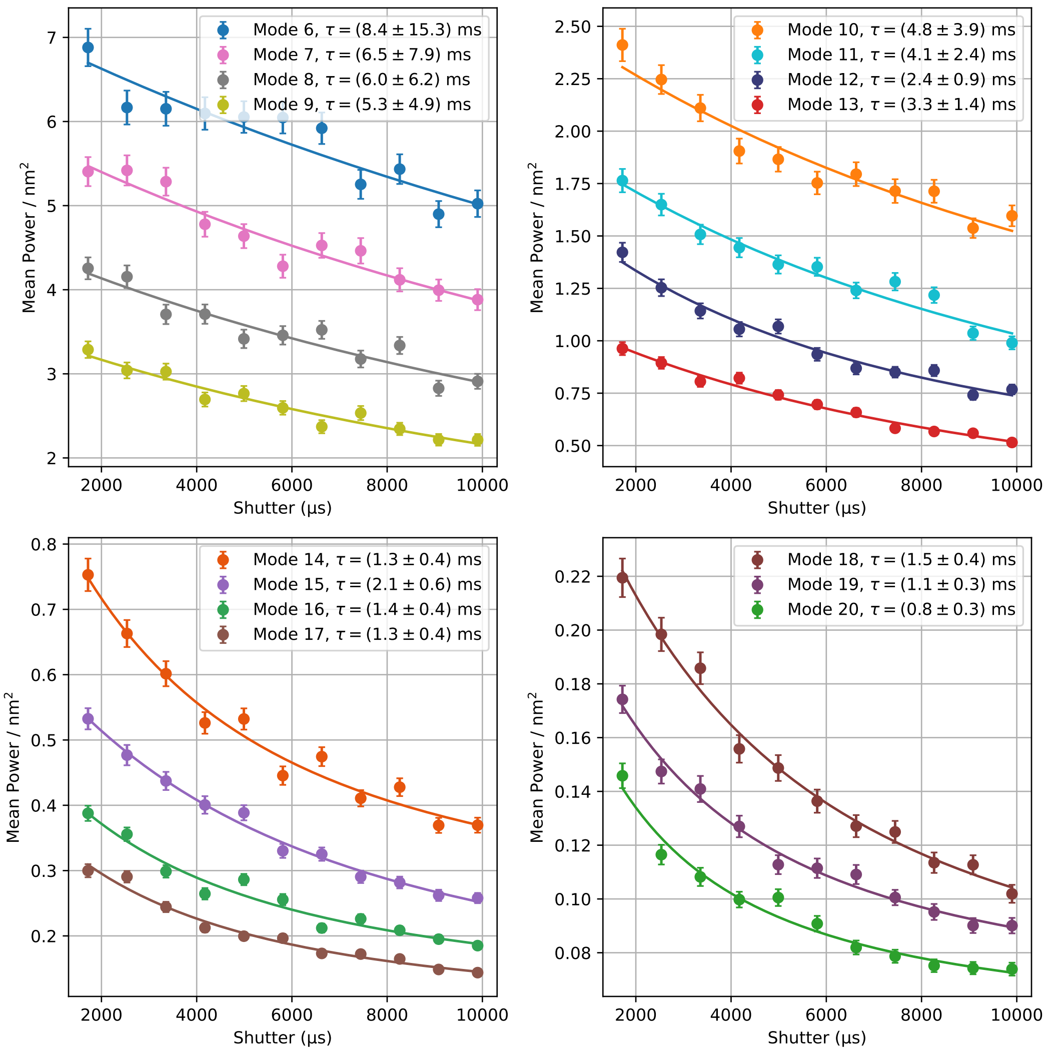


Figure S8: Power observed in all modes of a single cell at different exposure times shows the expected attenuation due to in-camera averaging. Fits using Eq. 2 determine the decay time of each mode. These are all modes for the cell used in Figure 11

#### REFERENCES


1. Yoon, Y.-Z., H. Hong, A. Brown, D. C. Kim, D. J. Kang, V. L. Lew, and P. Cicuta, 2009. Flickering Analysis of Erythrocyte Mechanical Properties: Dependence on Oxygenation Level, Cell Shape, and Hydration Level. *Biophys J* 97:1606–1615.

2. Pécréaux, J., H.-G. Döbereiner, J. Prost, J.-F. Joanny, and P. Bassereau, 2004. Refined Contour Analysis of Giant Unilamellar Vesicles. *Eur. Phys. J. E* 13:277–290.

3. Rautu, S. A., D. Orsi, L. Di Michele, G. Rowlands, P. Cicuta, and M. S. Turner, 2017. The Role of Optical Projection in the Analysis of Membrane Fluctuations. *Soft Matter* 13:3480–3483.

4. Kariuki, S. N., A. Marin-Menendez, V. Introini, B. J. Ravenhill, Y.-C. Lin, A. Macharia, J. Makale, M. Tendwa, W. Nyamu, J. Kotar, M. Carrasquilla, J. A. Rowe, K. Rockett, D. Kwiatkowski, M. P. Weekes, P. Cicuta, T. N. Williams, and J. C. Rayner, 2020. Red Blood Cell Tension Protects against Severe Malaria in the Dantu Blood Group. *Nature* 585:579–583.
